# Influence of attention mechanisms on cerebellar and basal ganglia activity during vocal emotion decoding

**DOI:** 10.1101/2024.11.20.624507

**Authors:** Leonardo Ceravolo, Marine Thomasson, Ioana Medeleine Constantin, Emma Stiennon, Émilie Chassot, Jordan Pierce, Alexandre Cionca, Didier Grandjean, Lukas Sveikata, Frédéric Assal, Julie Péron

**Affiliations:** Neuroscience of Emotion and Affective Dynamics Laboratory, Department of Psychology and Swiss Centre for Affective Sciences, University of Geneva, Switzerland; Cognitive Neurology Unit, Department of Neurology, University Hospitals of Geneva, Geneva, Switzerland; School of Physical and Occupational Therapy, McGill University, Montréal, QC, Canada; Clinical and Experimental Neuropsychology Laboratory, Department of Psychology and Swiss Centre for Affective Sciences, University of Geneva, Switzerland; Cognitive and Affective Neuroscience Lab, Department of Psychology, University of Nebraska-Lincoln, USA; Medical Image Processing Laboratory, Neuro-X Institute, École Polytechnique Fédérale De Lausanne (EPFL), Geneva, Switzerland; Faculty of Medicine, University of Geneva, Switzerland

## Abstract

Emotional prosody processing involves a widespread network of brain regions, but the specific roles of the cerebellum and basal ganglia in explicit and implicit tasks are not well known or understood. This study investigated how the cerebellum and basal ganglia contribute to explicit (emotion categorization) and implicit (gender categorization) processing of emotional prosody, namely when attention is directly versus implicitly oriented towards the emotion of the voice stimuli, respectively. Twenty-eight healthy French-speaking participants (average age: 65 years old) underwent high-resolution functional MRI while performing explicit and implicit vocal emotion processing tasks. Neuroimaging results revealed—and replicated—that both tasks recruited a widespread network, including the superior temporal cortex, inferior frontal cortex, primary motor and somatosensory cortices, basal ganglia, and cerebellum. The explicit task elicited stronger activations in the basal ganglia (caudate nucleus, putamen) and cerebellar regions (Crus I/II, lobules VI, VIIb, and X), consistent with higher cognitive control demands. In contrast, the implicit task was associated with activations in cerebellar lobules IV-V, VI, VIII, and IX, along with the thalamus. Regression-based functional connectivity analyses further demonstrated stronger connectivity between the right cerebellar lobule IX and the putamen, as well as the cerebellar vermis (XII), particularly during implicit processing. These findings highlight the distinct contributions of the cerebellum and basal ganglia to emotional prosody processing, with explicit tasks engaging associative and cognitive control networks, while implicit tasks rely more on sensorimotor and automatic neural processing mechanisms.

## Introduction

The cerebellum is increasingly recognized for its pivotal role in modulating both cognitive (Ito, 2008; Koziol et al., 2014) and emotional processes (Adamaszek et al., 2017), including the recognition of vocal emotional cues (Ceravolo et al., 2021; Pierce & Péron, 2020; Pierce et al., 2023; Thomasson et al., 2021; Thomasson et al., 2023), commonly referred to as ‘emotional prosody’ (Bänziger & Scherer, 2005; Grandjean, 2021; Grandjean et al., 2006). Emotional prosody encompasses the tone, rhythm, and melody of speech that convey emotional content, which is decoded by listeners through both explicit and implicit attentional focus. Substantial progress has been made in understanding the involvement of the cerebellum in emotional prosody recognition, particularly through its connections with the cerebral neocortex and subcortical structures such as the basal ganglia (Adamaszek et al., 2017; Bostan & Strick, 2018; Ceravolo et al., 2021; Pierce & Péron, 2020). Recent studies suggest that the cerebellum is part of a broader network that includes the basal ganglia, playing a critical role in processing emotions. The basal ganglia are notably involved in the rhythmic aspects of speech decoding and the regulation of emotional expression (Péron et al., 2013; Thomasson et al., 2022). However, much remains to be understood concerning topdown attentional mechanisms influencing cerebellar activity during this process (Ivry & Keele, 1989; Leiner et al., 1991; Schmahmann, 1997).

Attentional mechanisms are crucial for modulating emotional prosody processing (Ceravolo et al., 2016b; Frühholz et al., 2012; Grandjean, 2021; Grandjean et al., 2005; Pinheiro et al., 2019; Sander et al., 2005; Schirmer & Kotz, 2006). These mechanisms involve explicit attention, where participants deliberately focus on the emotional content of speech, or implicit attention, where the emotional tone is automatically processed while attention is directed elsewhere—for instance on speaker gender or voice spatial location. Recent research has shown that the response of the cerebellum to emotional prosody is influenced by these attentional mechanisms, but the precise role of the cerebellum in attention shifts during emotional decoding remains unclear (Frühholz et al., 2012; Stirnimann et al., 2018; Thomasson & Péron, 2022). Frühholz et al. (2012) demonstrated that explicit processing of emotional prosody activates different brain regions compared to implicit processing, emphasizing the importance of attention in emotional recognition. In this study, however, the role of the basal ganglia and the cerebellum was not specifically studied.

A recent meta-analytic study examined how attention allocation impacts cerebellar involvement in emotion processing in healthy participants (Pierce et al., 2023). This analysis included activation foci from 80 human neuroimaging studies of emotion, collectively covering 2761 participants. Results from explicit emotion processing tasks revealed clusters within the posterior cerebellar hemispheres (bilateral lobule VI/Crus I/II, the vermis, and left lobule V/VI), which were consistently activated across studies. In contrast, implicit emotion processing tasks activated clusters including bilateral lobules VI/Crus I/II, right Crus II/lobule VIII, anterior lobule VI, and lobules I-IV/V. A direct comparison between the two categories revealed five overlapping clusters in the right lobules VI/Crus I/Crus II and left lobules V/VI/Crus I of the cerebellum, suggesting involvement in both explicit and implicit emotion processing. Interestingly, explicit tasks showed greater activation in right lobule VI and left lobule VI/vermis, while implicit tasks activated left lobule VI more significantly. These findings support and extend previous research, indicating that posterior cerebellar regions contribute to both explicit and implicit emotion processing and are linked to neocortical networks involved in cognitive/executive functions, mentalizing, and salience processing.

Recent studies on the non-motor roles of the basal ganglia and cerebellum provide a broader perspective on their involvement in emotion processing (Pierce & Péron, 2020; Schutter & Van Honk, 2005; Strata, 2015). Theoretical models suggest that organisms build internal models to predict environmental changes, allowing for adaptive responses before changes occur (Koziol et al., 2014). The cerebellum is particularly involved in predictive adaptation concerning temporal structures (Cabaraux et al., 2020; Ito, 2008). Over time, experiences are grouped into functional units known as “chunks” (Graybiel, 1998), facilitating automatic and ‘habitual’ responses. This shift from goal-directed processing to more habitual modes is supported by transitions between associative-limbic and sensorimotor pathways in the cerebellum and basal ganglia (Graybiel, 2008; Habas et al., 2009; Krack et al., 2010). In the domain of emotion and affective neuroscience, this may explain why certain vocal patterns conveying emotions transition from conscious control to automatic processing—potentially mediated by the cerebellum, though empirical studies in this area remain limited.

The present study aims to build on these findings by investigating how the modulation of attentional focus may affect the involvement of the cerebellum and basal ganglia in emotional prosody decoding. In light of above-mentioned literature and considering the current state of knowledge in the field, our study seeks to address the following primary research question: What is the impact of attentional focus mechanisms on the involvement of cerebellar and basal ganglia brain regions in emotional (anger-happiness-neutral) prosody processing? Using functional magnetic resonance imaging (fMRI), we examined how changes in attentional focus—explicit versus implicit vocal emotion processing—would affect cerebellar and basal ganglia activity in healthy older adults. By employing an interdisciplinary approach that combines experimental psychology with neuroimaging, we aimed to deepen our understanding of the roles of the cerebellum and basal ganglia in emotion processing within these networks. Based on above-mentioned research by us and others, we therefore hypothesized that both the basal ganglia and cerebellum would not only be involved in emotional prosody recognition but also that their activity would be modulated by attentional demands, with greater engagement of the associative parts of these brain structures under explicit processing conditions (medial putamen, caudate, globus pallidus, cerebellar lobules V/VI, Crus I/II, vermis; Pierce and Péron (2020); Pierce et al. (2023)) and of their sensorimotor parts under implicit processing conditions (globus pallidus, putamen, cerebellar Crus I/II and lobules I-V, VI, VIII; Ceravolo et al. (2021); Pierce and Péron (2020); Pierce et al. (2023)). Furthermore, we predicted that this activity pattern would be stronger for emotional (angry, happy) vs. neutral voice prosody. Based on structural and functional connectivity research in the field (Ceravolo et al., 2021; Pierce & Péron, 2020), we also expected enhanced functional connectivity between these regions of the basal ganglia and cerebellum for each task.

## Materials and Methods

### Participants

Twenty-eight healthy participants were recruited for the study (13 female or 46.52%, 15 male or 53.48%). They were all French speakers with a mean age of 65.4 years (SE=8.9, range=48-77). Older adults were recruited as age-matched controls as part of a study of stroke patients that will be described elsewhere in the future. According to the Edinburgh Handedness Inventory criteria (Oldfield, 1971), 26 participants were right-handed and 2 were left-handed. Their mean education level was 16.4 years (SE = 3.0, range = 12-22). Exclusion criteria for the recruitment were: 1) history of neurological disorders, 2) head trauma, 3) anoxia, stroke or major cognitive deterioration, as attested by their score on the Montreal Cognitive Assessment (Nasreddine & Patel, 2016) (mean score = 28, SD = 1.3, range = 25-30). Moreover, none of them wore hearing aids or had a history of tinnitus or a hearing impairment, as attested either by their PEGA (‘Protocole d’Evaluation des Gnosies Auditives’) score (mean = 28.7, SD = 1.4, range = 25-30). All participants gave their written informed consent and the study was approved by the local ethics committee. Using a two-tailed testing method with a medium effect size of 0.5, an alpha of 0.05 and our sample of N=28, we obtain a posthoc power of 72.27% as calculated in G*Power 3.1.9.7 (Faul et al., 2007) for comparing our tasks (mean comparison in one sample) taking into consideration the Emotion factor.

### Vocal emotion recognition task procedure

The vocal stimuli consisted of two speech-like but semantically meaningless sentences extracted from Banse and Scherer’s validated database (Banse & Scherer, 1996). These pseudo-sentences were spoken in a neutral—more specifically the “interest” emotion category in the database, happy (‘elation’) or angry (‘hot anger’) tone by two male and two female actors, resulting in a total of 24 different stimuli. Among these 24 stimuli, 18 were used in the present study and assigned to each task and block (see Fig.1A-C), keeping the balance between actor gender and emotions. During scanning, these binaurally recorded auditory stimuli were played through MRI-compatible headphones and displayed using E-Prime (http://www.pstnet.com/eprime.cfm) in two distinct runs (Fig.1A). Participants performed a categorization task with two levels of the Task factor (implicit, explicit) and three for the Emotion factor (anger, happiness, neutral). See Fig.1B for task details and illustration. This paradigm was divided into 16 counterbalanced blocks (6 trials per block), for a total of 96 trials per run and a grand total of 192 trials per participant. For four of these blocks (implicit), participants had to make a gender decision (participants had to indicate “male” or “female” by pressing a key with their index or middle finger). For the other four blocks (explicit), participants performed an emotion categorization task (they had to indicate “happiness” or “neutral” / “anger” or “neutral” by pressing a key with their index or middle finger). For explicit blocks, the participants did not know in advance whether they would be presented with happy and neutral or angry and neutral trial types. For all blocks, the order of the emotion presented was pseudo-randomized to avoid repetitions of the same emotion several times in a row and to avoid boundary effects (alternating emotion as the last trial of each block).

**Fig.1:**
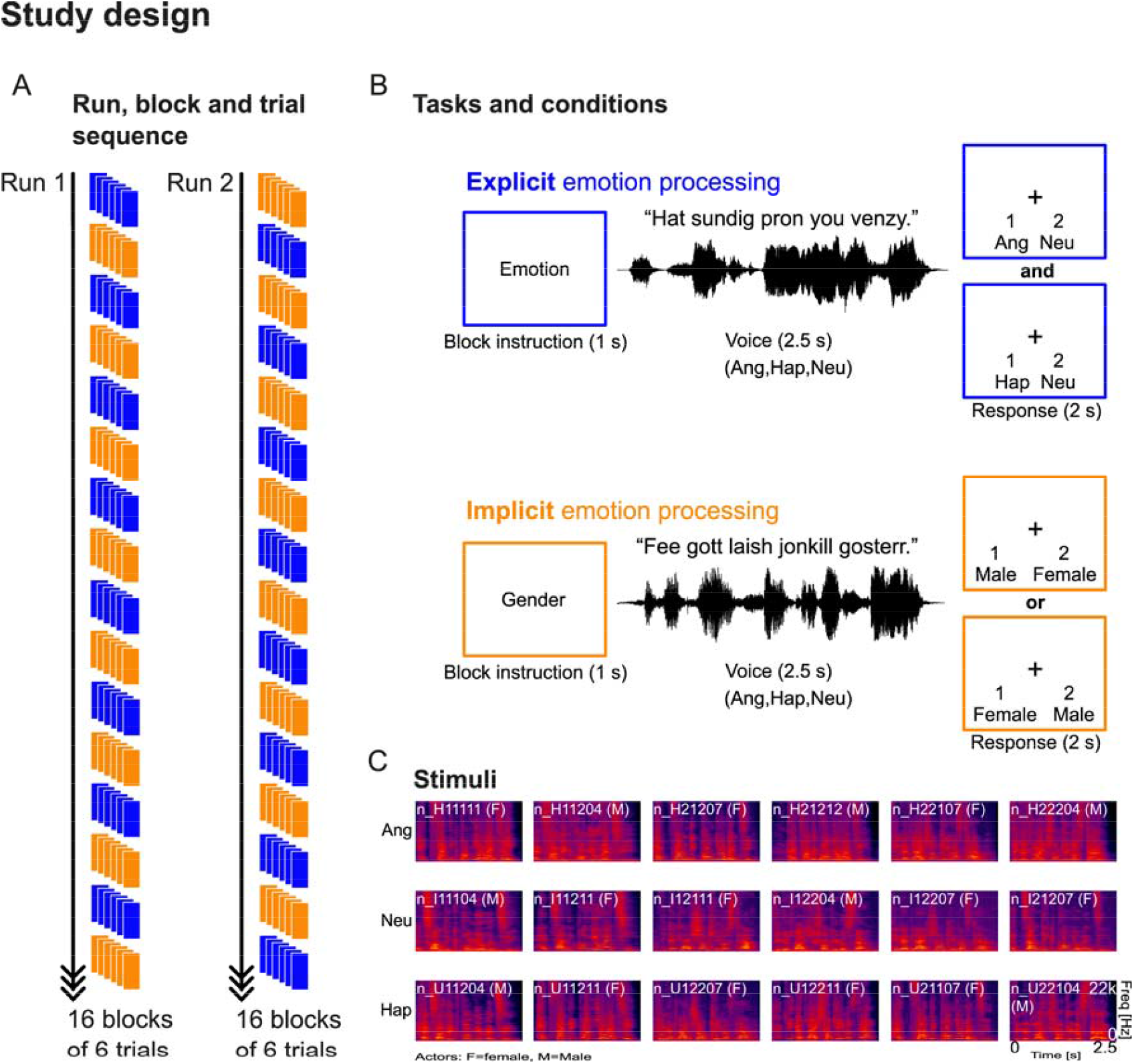
Study design, task and stimuli. ‘A’ Tasks were split into two runs of equal length including 16 blocks of 6 trials each, alternating between the explicit and the implicit vocal emotion processing tasks (blue and orange, respectively). These tasks are described in ‘B’, with first an instruction of the type of block to come, then the stimulus is presented for 2.5 s and the participants had to categorize the stimulus according to the instructed task (2 s). ‘C’ Illustration through a spectrogram of the 18 stimuli used in the study, with stimulus identity and speaker gender in parentheses (F: female, M: male). Ang: anger; Hap: happiness; Neu: neutral.

We assumed that during the gender decision task, participants would focus on the gender of the voice, but the emotional tone of the voice still would be processed implicitly—which was the aim of this task, namely an implicit emotion processing. Before each block, a message indicating “gender” or “emotion” was displayed on the screen for 1 sec as an instruction for the coming block. The vocal stimuli were then played (duration: 2.5 s; Fig.1C) and a fixation cross was presented at the center of the screen, and the two possible responses displayed on the screen for 2 s. Each of the two runs contained 96 trials and lasted approximately 10 minutes.

### Behavioral data analysis

Behavioral results illustrate the probability of accurately recognizing voice gender or emotion (Task factor) as a function of emotional tone (Emotion factor). These analyses were computed using mixed effects regression analyses for the response accuracy (of-interest) and for the associated reaction times (of-no-interest) in R studio (Allaire, 2012).

For the ‘response’ dependent variable, we used a logistic regression using the lme4 ‘glmer’ package (Bates et al., 2015) with Emotion (Emo) interacting with Task and Run as fixed effects, the random slope of stimulus identity (Stimuli) and Task as a function of participant identity (ParticipantID; uncorrelated). Furthermore, participants’ age (ParticipantAge), Gender (ParticipantGender), stimuli order (StimuliOrder), voice actor identity (ActorID) and Gender (ActorGender) were included as random factors. The model formula was the following:

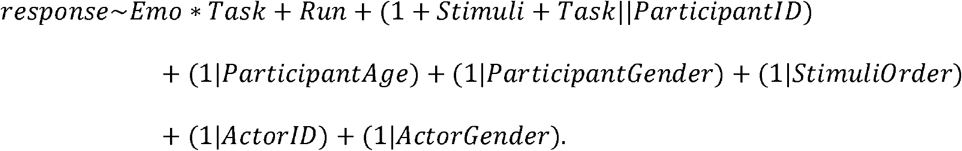

For the ‘reaction times’ dependent variable, we used a linear regression using the Ime4 ‘lmer’ package with the exact same formula as above.

For our planned comparisons, we used a confidence interval of 97.5% and a Bonferroni correction for multiple comparisons. Specific contrasts were computed using the ‘phia’ package (De Rosario-Martinez et al., 2015).

No difference was observed between models including or excluding participant gender as a random effect (AIC_geder_=3121.1 vs. AIC_no_-_geder_=3119.1, p>.1).

### MRI data acquisition

All functional imaging data were recorded on a 3-T Siemens Trio System (Siemens, Erlangen, Germany) scanner equipped with a 32-channel antenna. A 3D sequence was used to acquire a high-resolution T1-weighted image (1mm isotropic voxels; TR=1900msec; TE=2.27ms; matrix resolution=256 x 256; FA=9 degrees; 192 total slices).

For the task, functional images were acquired in descending order using a multi-band echo-planar imaging (EPI) sequence of 54 slices aligned along the anterior-posterior commissure (voxel size: 2.5mm isotropic; TR=1300msec; TE=20ms; FOV=205 x 205mm; matrix resolution=84 x 84; FA=64 degrees; BW=1952Hz/px). Functional image acquisition was continuous throughout each run of the task.

### MRI data analysis

Data were preprocessed using a mix of SPM12 (SPM12, Wellcome Trust Centre for Neuroimaging, London, UK), the CONN toolbox (Whitfield-Gabrieli & Nieto-Castanon, 2012) and the Artifact detection tools (ART) toolbox (https://www.nitrc.org/projects/artifact_detect/). Functional data were first converted to 4D Nifti in SPM12 for each run separately to get one single file per run, for more efficient computation performance and decrease storage volume. Preprocessing steps included the following: realignment and unwarping, slice-timing correction, artifact detection followed by an additional denoising (scrubbing including functional regression and functional bandpass [0.01-0.1Hz]). At this stage, direct segmentation and normalization into the Montreal Neurological Institute (MNI) space (Collins et al., 1994) was performed for improved comparison between the brain tissues of our sample of participants and other existing fMRI studies. Finally, data were spatially smoothed with an isotropic Gaussian filter of 8 mm full width at half maximum. Following an initial step consisting in the creation of a participant-specific matrix including the onset and reaction times of each trial of each run, preprocessed, smoothed functional images were then analyzed with SPM12 both at the participant-level (‘first-level’) as well as at the group-level (‘second-level’). A single general linear model was used to compute first-level statistics, using a convolution with the Hemodynamic Response Function (HRF). Each event was therefore modelled using HRF function, locked to the onset of each voice stimulus using continuous MRI data acquisition. The design matrix included both runs separately. Within each run, columns represent the Emotion factor for each task (Task factor): for the implicit task there are three columns for each emotion (anger, happiness, neutral) and four columns for the explicit task considering each association (anger, neutral; happiness, neutral). In addition, six motion parameters were included as regressors of no interest to account for movement in the data, as well as trial-level reaction times since only response accuracy was of interest. Error trials were also included as regressors of no interest per run. This error regressor also included trials with reaction times shorter than 200 ms since it was highly unlikely that the participants could accurately respond this fast. In average, this ‘Error regressor’ contained 9.68 (SD=2.64) trials per participant out of the total 196 trials, namely 4.37% of the total count. Regressors of interest were used to compute simple contrasts for each participant, leading to separate main effects of Angry, Neutral and Happy voices for each task, for a total of six simple contrasts (Implicit task: Angry, Neutral and Happy voices; Explicit task: Angry, Neutral and Happy voices).

This model yielded two flexible factorial second-level analyses, with factors Task and Emotion. We also had the Participants factor (Factor 1 in each analysis) to consider interindividual variability, with data independence set to ‘yes’, variance to ‘unequal’ while for the within factors of interest data independence set to ‘no’, variance to ‘unequal’. Second-level models focused on: the Task factor (Model 1) and on the interaction between Task and Emotion (Model 2, Task * Emotion).

Activations were ultimately thresholded in SPM12 by using a voxel-wise FDR correction at *p*<.05. Thresholded contrast activations were then rendered on brains from the CONN toolbox (Whitfield-Gabrieli & Nieto-Castanon, 2012). Regions were labelled using the latest version of the ‘automated anatomical labelling’ (‘aal3’) atlas (Rolls et al., 2020; Tzourio-Mazoyer et al., 2002).

### Functional connectivity analyses

To test for un-mediated connectivity in both tasks and between our regions of interest (ROIs; N=32), namely all subregions of the basal ganglia and cerebellum, we computed undirected functional connectivity analyses using task-based, generalized psychophysiological interactions (gPPI) with bivariate regression in CONN (Whitfield-Gabrieli & Nieto-Castanon, 2012). The 32 ROIs correspond to all basal ganglia, thalamus and cerebellar regions of the ‘aal3’ atlas implemented in the CONN toolbox (Rolls et al., 2020; Tzourio-Mazoyer et al., 2002). Paraphrasing the authors of the toolbox, “The gPPI measures represent task-modulated effective connectivity between ROIs, namely changes in association strength covarying with an experimental factor. They are aimed at investigating task-related modulation of connectivity patterns in event-related task designs”. These analyses are computed using a separate multiple regression model for each ROI, including as independent variables: 1) selected task effects convolved with the HRF (psychological factor); 2) the seed ROI signal across time (physiological factor); and 3) the interaction term between 1 and 2. The output of gPPI corresponds to the map of regression coefficients associated with the interaction term in these models. The results analyzed and presented in Fig.S1 are thresholded using parametric multivariate statistics with cluster-level inference, including a ROI-level MVPA omnibus test thresholded at *p*<.05 FDR to take into account multiple comparisons (Jafri et al., 2008). In our view, this analysis yields very robust task-based functional connectivity that shares some critical aspects with effective connectivity but lacks the direction of the causality between ROIs. We therefore categorize these results as ‘functional connectivity’.

## Results

This section reports the behavioral and neuroimaging results of our sample of participants (N=28), who were asked to categorize the gender (implicit emotion processing) or the emotion (explicit emotion processing) of angry, happy or neutral voices presented to them via headphones while lying in the fMRI scanner. The explicit and implicit tasks define the levels of the Task factor, while the emotion conveyed by each voice stimulus (anger, happiness, neutral) constitutes the levels of the Emotion factor. Main effects as well as partial and full interaction effects are described below.

### Behavioral data

Behavioral data included one dependent variable of interest—response accuracy (Fig.2)— and one of no-interest, namely reaction times (see supplementary material). Response accuracy results are presented below and summarized in Table S1—including reaction times.

**Fig.2:**
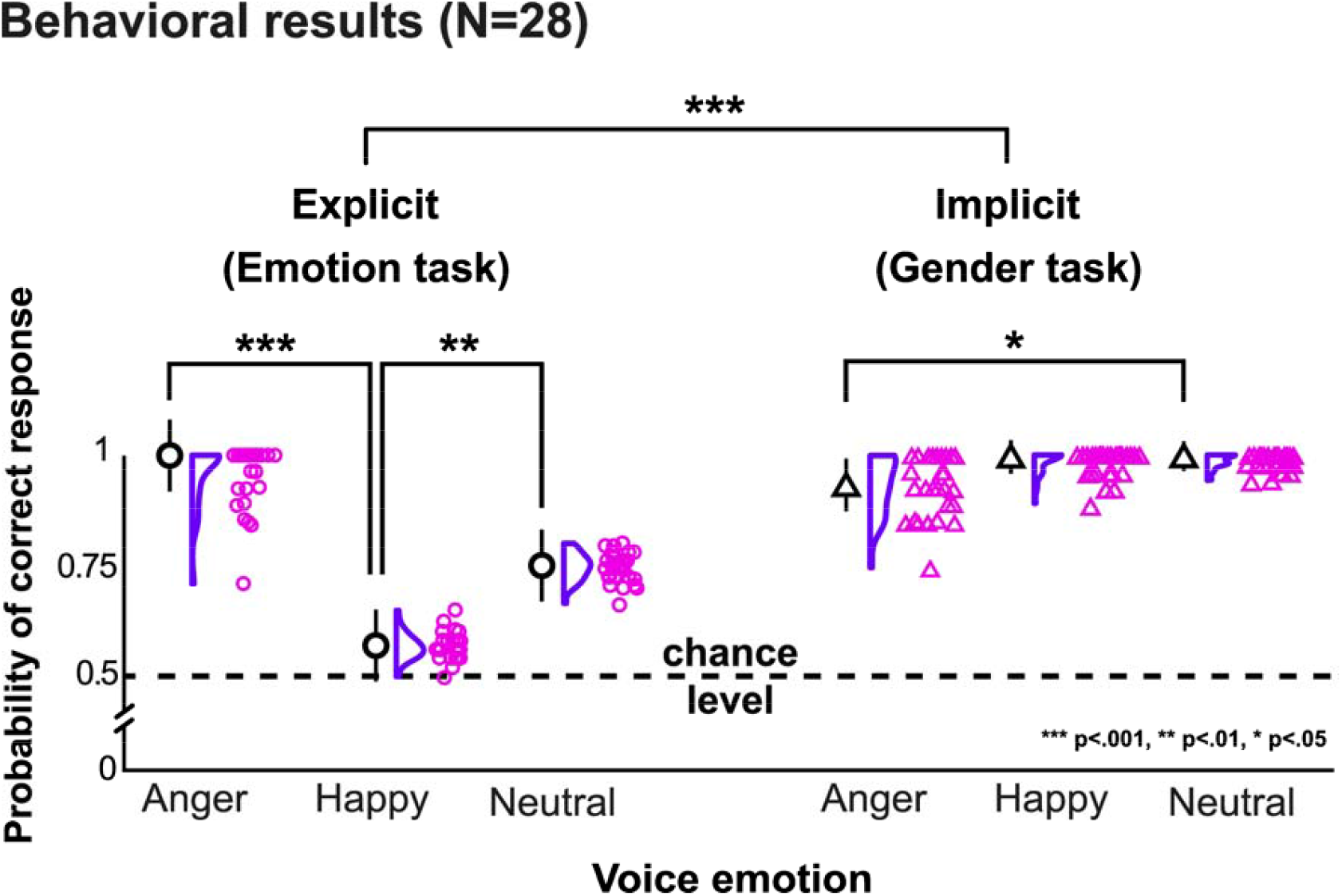
Behavioral results for response probability in the Gender (Implicit) and the Emotion (Explicit) task. Circles and triangles represent individual values for each participant (N=28; circles: Emotion task, triangles: Gender task), with data curves as half-violin distribution and mean values with error bars representing the standard error of the mean (Y axis) as a function of voice emotion (X axis). Chance level for response probability is 50% (0.5, Y axis) since all tasks involved a 2-alternate forced choice type of response. *** *p*<.001, ** *p* <.01, * *p* <.05. Descriptive values are reported in Table SI.

#### Response patterns and accuracy

Response data of each task were treated as ‘correct’ or ‘incorrect’ and coded as ‘1’ or ‘0’, respectively. Logistic regression was used to better characterize data and allow for the interpretation of the probability of a correct response as a function of our Task and Emotion factors (fixed effects)—including also the split into two runs and taking into consideration relevant random effects (see Methods).

The probability of a correct response was predicted by the Task factor with higher accuracy for the gender task (χ^2^(1)=93.33, *p*<.001), voice Emotion (χ^2^(2)=19,11, *p* <.001), Run (with higher accuracy in run 2, χ^2^(1)=8.81, *p* <.01) and the interaction between Task and Emotion (χ^2^(2)=85.73, *p* <.001). Across tasks, neutral and angry voices yielded a higher probability of correct response compared to happy voices for the Emotion factor (χ^2^(1)=5.54, *p* <.05). The interaction between Task and Emotion was mainly explained by an inverted pattern of response between tasks, in which higher accuracy probability was observed for angry compared to neutral and happy voices in the explicit task. The Task * Emotion interaction also led to higher accuracy in the gender task for angry voices ([Task Gender > Task Emotion * Angry > Neutral + Happy voices]: χ^2^(1)=84.88, *p* <.001). Additionally, differences were observed in accurate response probability between angry vs happy voices in the Emotion task (χ^2^(1)=32.59, *p* <.001)—but not in the Gender task (χ^2^(1)=1.74, *p* =.19)—and also between tasks, with higher accuracy for angry vs happy in the Gender task as opposed to the inverse in the Emotion task (χ^2^(1)=71.97, *p*<.001). Response probability for angry vs neutral voices also differed in both tasks (Gender: χ^2^(1)=6.49, *p* <.05; Emotion: χ^2^(1)=9.23, *p* <.01) and between tasks (χ^2^(1)=53.46, *p* <.001) as well as for happy vs neutral voices in the Emotion task only (χ^2^(1)=8.99, *p* <.01; Gender task: χ^2^(1)=1.21, *p* >.10; Gender vs Emotion task: χ^2^(1)=3.15, *p* <.10). Variance explained by the fixed effects was 38.24% (R^2^c) while the full model including random effects (R^2^c) explained 61.67% of the variance of the response probability data. See Fig.2 for details and illustration.

### Neuroimaging data

Functional images were continuously acquired throughout each participant’s session to investigate brain correlates associated with the two tasks (explicit and implicit vocal emotion processing, Fig. 3), the emotional content when contrasting tasks (angry, happy, and neutral voices for explicit > implicit and implicit > explicit), and the interactions between task type and emotion. Whole-brain voxel-wise statistics are presented below with correction for multiple tests (false-discovery rate, ‘FDR’). Since reaction times were of no-interest and participants were not instructed to respond as fast as possible, reaction times were used as a trial-level covariate of no interest in the neuroimaging statistical models. Across tasks, whole brain correlates included a widespread network of activated brain regions in the superior temporal and inferior frontal cortex (Fig.3A,C), primary motor and somatosensory cortex (Fig.3A-F), as well as in the basal ganglia (Fig.3G) and cerebellum (Fig.3H,I).

**Fig.3:**
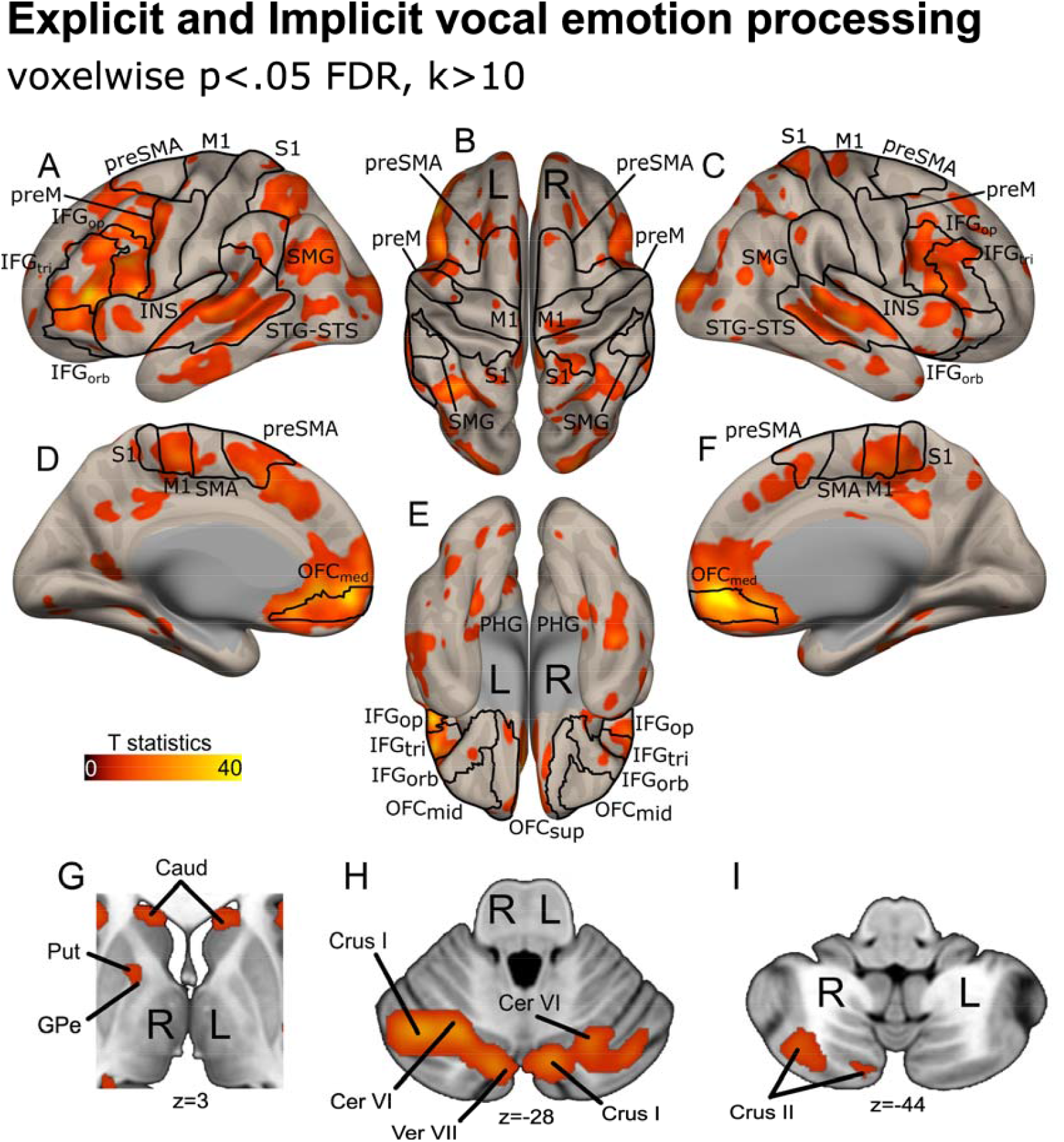
Whole-brain activations for explicit and implicit voice emotion processing. Lateral and medial cortical (lateral cortex ‘A-C’; medial cortex ‘D-F’) as well as basal ganglia (‘G’) and cerebellar activations (‘G-I’) across tasks. All activations reported using an FDR corrected voxel-wise threshold of *p* <.05. Color bars represent T-values. IFG: inferior frontal gyrus; INS: insula; STG: superior temporal gyrus; STS: superior temporal sulcus; preSMA: pre-supplementary motor area; SMA: supplementary motor area; preM: premotor cortex; PHG: parahippocampal gyrus; M1: primary motor cortex; S1: primary somatosensory cortex; OFC_med_; medial orbitofrontal cortex; SMG: supramarginal gyrus; Put: putamen; Caud: caudate nucleus; GPe: globus pallidus, extern portion; Cer: cerebellar lobule; Ver: vermis. Suffixes: tri, *pars triangularis;* op, *pars operations;* orb, *pars orbitalis;* med, medial; sup, superior; mid, middle. L: left hemisphere, R: right hemisphere.

#### Effects of Explicit vs. Implicit Processing

Both tasks were designed to highlight the levels of vocal emotion processing: focusing on voice gender or on expressed emotion to implicitly or explicitly process vocal emotion, respectively. We present here the results with a focus on the basal ganglia and cerebellum (see signal change in Fig.4H), while whole-brain data are visible in Fig.S2.

**Fig.4:**
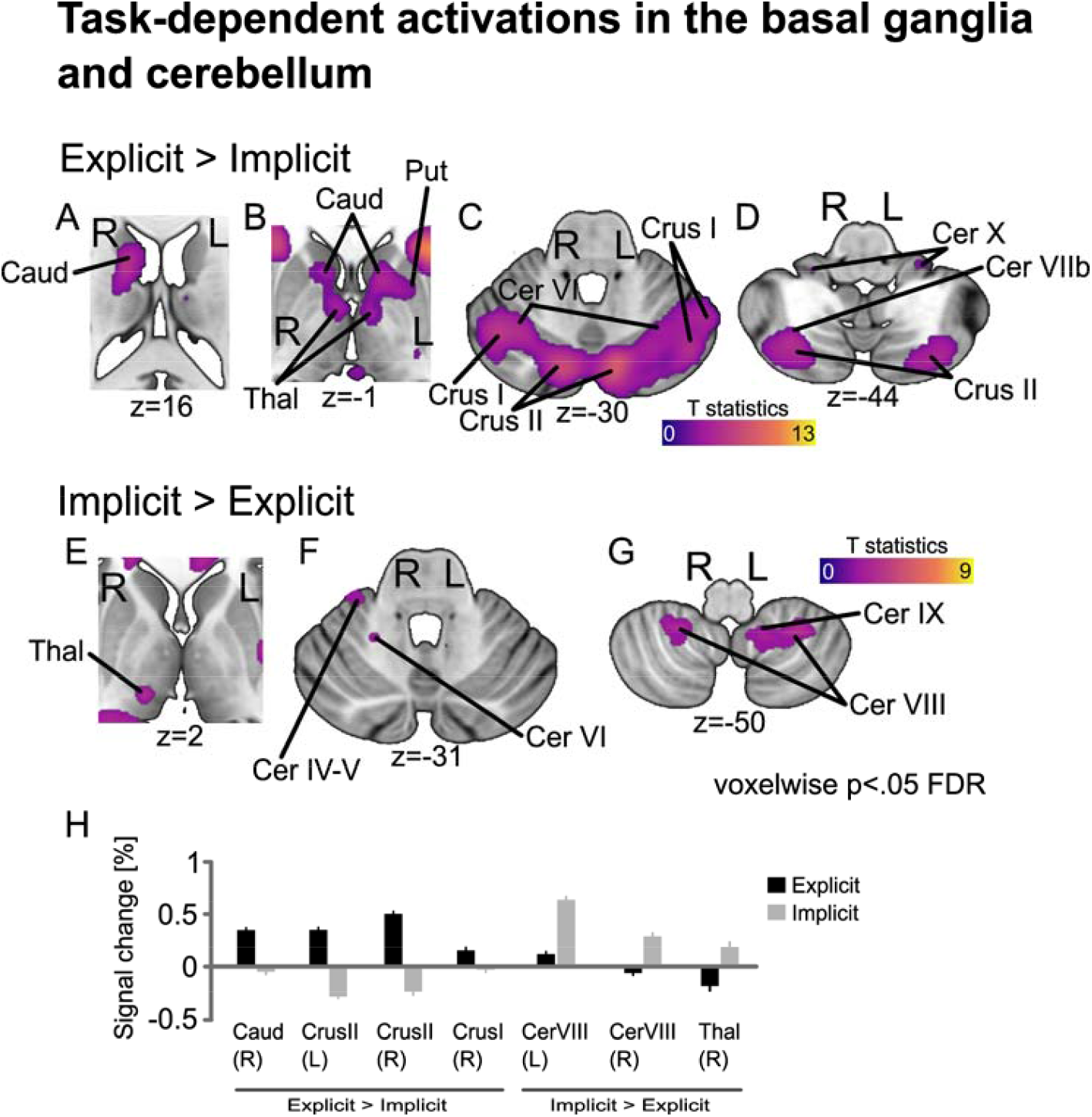
Cerebellum and basal ganglia activations for the Task contrasts. Above-threshold activations when contrasting explicit to implicit (‘A-D’) emotion processing tasks (Explicit>Implicit) and its inverse (‘E-G’; Implicit>Explicit). ‘H’ Percentage of signal change in regions of both contrasts, obtained by selecting the peak voxel for each participant and taking the nearest maxima, then using singular value decomposition to select among 27 contiguous voxels the ones that explained at least 80% of the variance in signal in the peak. Error bars represent the standard error of the mean. All activations reported using an FDR corrected voxel-wise threshold of *p* <.05. Color bars represent T-values. Caud: caudate nucleus; Put: putamen; Thal: thalamus; Cer: cerebellar lobule. L: left hemisphere, R: right hemisphere.

The contrast between explicit vs implicit processing of vocal emotions showed enhanced activations in the caudate nucleus, putamen, thalamus (Fig.4A,B), as well as in widely spread regions of the cerebellum including Crus I/II and cerebellar lobule VI (Fig.4C) and in lobules X and VIIb (Fig.4D). See also Table S2.

In the inverse contrast, implicit vs explicit vocal emotion processing, we observed distinct activations including the thalamus (Fig.4E), cerebellar lobules IV-V and VI (Fig.4F), and cerebellar lobules VIII and IX (Fig.4G). See Table S3.

#### Emotion-Specific Effects

*We* then contrasted the tasks and looked at each emotion individually in the cerebellum and basal ganglia. These results are presented in Fig.5 (Table S4-7). Whole-brain, cortical activations for these contrasts are illustrated in Fig.S3-S8.

**Fig.5:**
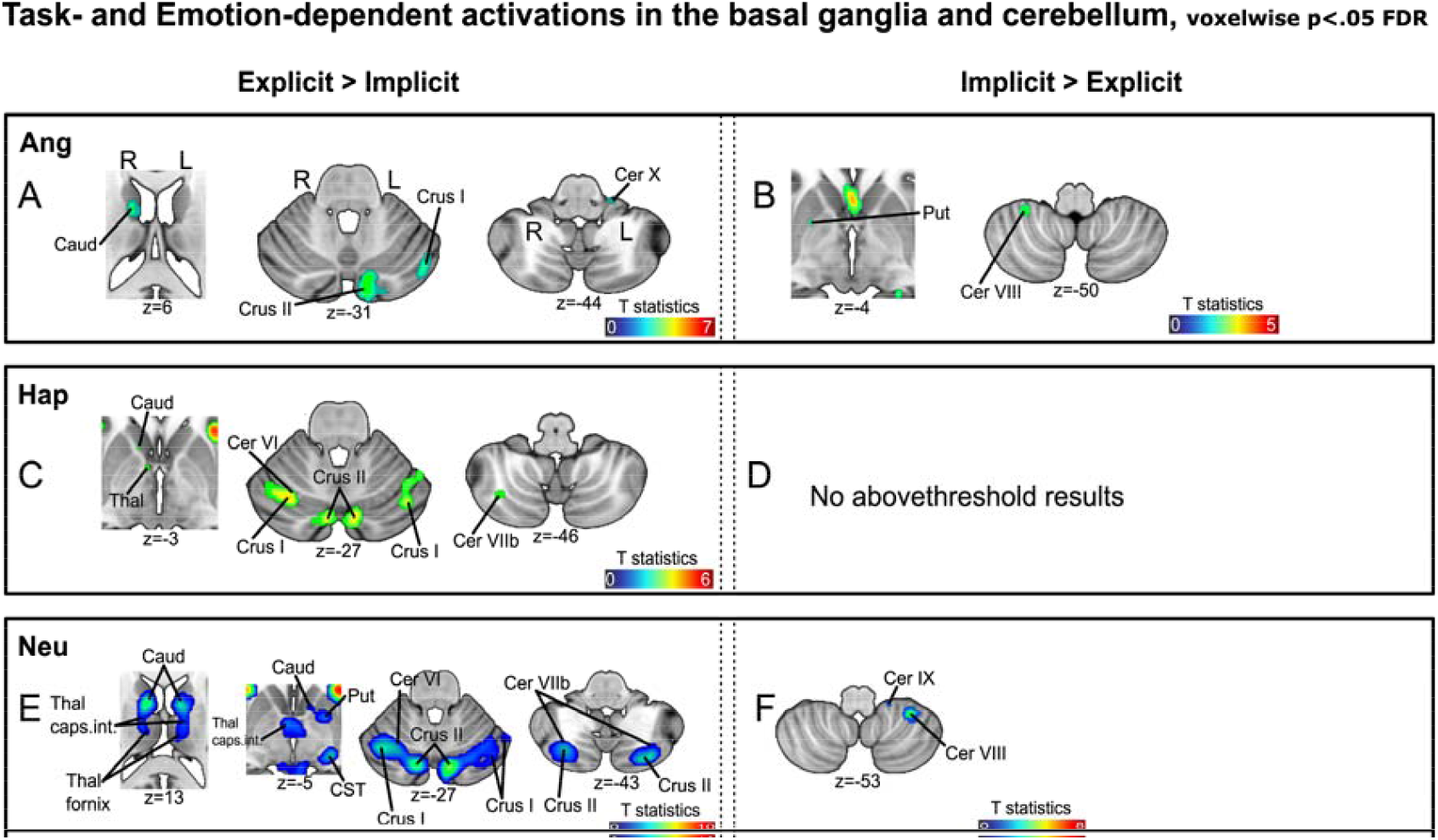
Basal ganglia and cerebellum activations contrasting the levels of the Task factor for each emotion. Basal ganglia and cerebellar activations when contrasting explicit to implicit tasks for angry (‘A’) happy (‘C’) and neutral voices (‘E’). Basal ganglia and cerebellar activations when contrasting explicit to implicit tasks for angry (‘B’) happy (‘D’) and neutral voices (‘F’). All activations reported using an FDR corrected voxel-wise threshold of *p*<.05. Color bars represent T-values. Caud: caudate nucleus; Put: putamen; Thal: thalamus; Thal caps.int.: capsula interna of the thalamus; CST: cortico-spinal tract of the brainstem; Cer: cerebellar lobule. L: left hemisphere, R: right hemisphere.

When contrasting Explicit to Implicit emotion processing, we observed an overlap for angry, happy and neutral voices, especially in the caudate nucleus and crus II area of the cerebellum (Fig.5A,C,E). For the inverse Implicit > Explicit contrast, there was not much overlap between emotions, with activations in the putamen and left cerebellar lobule VIII for angry voices (Fig.5B) and in right cerebellar lobules VIII and IX for neutral voices (Fig.5F) while no significant voxels in these regions were observed for implicitly processed happy voices.

Other results pertaining to the Task * Emotion interaction—happy vs neutral voices in both tasks as well as angry vs happy (and inverse) in the explicit task—are reported Tables S8-S11.

### Functional Connectivity

Across tasks (explicit and implicit vocal emotion processing), these analyses generally led to direct and positive connectivity between the right cerebellar lobule IX and left and right putamen and the vermis (XII). Very similar results were obtained for the implicit task while no results reached significance for the explicit task. See Fig.S1 for illustration.

## Discussion

The present study investigated the impact of attentional focus on the involvement of cerebellar and basal ganglia activity in emotional prosody processing, contrasting explicit and implicit tasks—in which the participants focused their attention on voice emotion or voice gender, respectively. Our behavioral and neuroimaging data underline the role of the cerebellum and basal ganglia in modulating emotional prosody perception depending on task demands and specific emotional content.

### Behavioral Findings

Behavioral results revealed that response accuracy was influenced by both Task and Emotion factors. Our participants exhibited higher accuracy in the implicit task compared to the explicit emotion recognition task, suggesting that gender recognition (with implicit emotion processing) may be less cognitively demanding or more automated. Furthermore, voice emotion modulated response accuracy, with neutral and angry voices yielding better performance compared to happy voices. This effect was especially pronounced in the explicit task, in which accuracy was higher for angry voices compared to neutral and happy voices, suggesting that anger may be more salient and easier to categorize in an explicit emotional context.

The interaction between Task and Emotion further revealed an inversion in response patterns, with higher accuracy for angry voices in the explicit task but for neutral voices in the gender task. These findings suggest that attentional focus modulates the ease with which different emotional prosodies are processed, potentially due to differences in salience or cognitive demands associated with recognizing emotions versus categorizing gender from vocal cues. These data are consistent with research on explicit vocal emotion processing and recognition (Scherer, 1995) but also with data from emotional cueing paradigms by vocal emotion—in which voice signals are processed in an implicit manner—or in implicit vocal emotion processing paradigms (Brosch et al., 2008, 2009; Ceravolo et al., 2016a, 2016b; Frühholz et al., 2012; Grandjean et al., 2005; Sander et al., 2005). Study results, however, may vary depending on whether positive emotions are included in the design or if only negative emotions are used as stimuli (Bach et al., 2008).

### Neuroimaging Findings

Our fMRI results confirmed the involvement of a widespread network during vocal emotion processing, encompassing the superior temporal cortex, inferior frontal cortex, primary motor and somatosensory cortex, but more importantly for the scope of the present article: the basal ganglia and the cerebellum. These results align perfectly with previous studies highlighting the role of these regions in voice prosody perception and emotional salience processing (Frühholz et al., 2012; Grandjean, 2021; Pierce & Péron, 2020; Pierce et al., 2023). We will now discuss task-related effects in the next section.

#### Effects of Explicit vs. Implicit Processing

When contrasting explicit to implicit emotion processing tasks, we observed activations in the basal ganglia (bilateral caudate nucleus, left putamen), bilateral thalamus, and the cerebellum (bilateral Crus I/II, lobules VI and X, and right lobule VIIb). These findings suggest that explicit emotion recognition recruits additional resources related to cognitive control, salience detection, and motor coordination (Adamaszek et al., 2017; Corbetta & Shulman, 2002; Ito, 2008; Leiner et al., 1991; Middleton & Strick, 2000; Pierce et al., 2023). Conversely, implicit processing (implicit vs explicit contrast) was associated with activations in cerebellar lobules IV-V, VI, VIII, and IX, along with the right thalamus, suggesting a role for these regions in more automatic and sensory-driven aspects of emotional prosody perception (Ceravolo et al., 2021; Péron et al., 2013; Pierce & Péron, 2020).

Interestingly, functional connectivity analyses revealed robust, direct task-based connectivity between the right cerebellar lobule IX and both the left and right putamen, as well as the cerebellar vermis (XII). This connectivity pattern was more pronounced in the implicit task, suggesting a potential role for these regions in supporting non-conscious, automatic emotional decoding processes. This interpretation is in line with our previous research using implicit vocal emotion processing (Ceravolo et al., 2021; Pierce et al., 2023). In contrast, explicit emotion processing did not yield significant connectivity effects, potentially indicating a shift towards more cortical networks involved in conscious appraisal, including the amygdala, superior temporal lobe and the inferior frontal gyrus (Frühholz et al., 2012).

#### Emotion-Specific Effects

Examining task-related activations for specific vocal emotions further elucidated the underlying role of the cerebellum and basal ganglia. The explicit versus implicit contrast revealed overlapping activations in the caudate nucleus and Crus II of the cerebellum across all emotions (angry, happy, neutral), emphasizing a generalized role for these structures in conscious emotion recognition, as mentioned in the previous section. However, in the implicit versus explicit contrast, activation patterns differed between emotions, with putamen and left cerebellar lobule VIII involvement for angry voices and right cerebellar lobules VIII and IX for neutral voices, while happy voices did not show comparable effects. These results suggest that the implicit processing of emotional prosody relies on distinct subcortical and cerebellar networks depending on the emotional content, possibly reflecting differences in salience, familiarity, or motor simulation demands. These data are consistent with previous research on implicit vocal emotion processing, although the absence of results for happy voices was somewhat unexpected. On the contrary, implicitly processing happy voices usually leads to enhanced activity in the cerebellum and/or basal ganglia (Ceravolo et al., 2021). This discrepancy could be attributed to the task itself, or to the specific voice stimuli used—although these were highly validated—and it would require further investigation in future work.

### Limitations

While our study provides valuable insights into the interplay between attentional focus, cerebellar-basal ganglia connectivity, and emotional prosody processing, several limitations should be considered. First, our sample was limited to healthy older adults, making it difficult to generalize findings to younger populations or clinical groups with cerebellar or basal ganglia dysfunctions. As mentioned in the methods, part of this dataset represents matched-control participants for a comparison with stroke patients that will be presented elsewhere in the future. Second, the study design focused on categorical emotion classification rather than on a more dimensional approach, which may limit the interpretation regarding the full spectrum of affective processing. Third, even though the tasks both involved a 2-alternate forced choice paradigm—an adaptation was made for the explicit task, see methods—it is still difficult to ignore the fact that the Emotion factor yielded differences between tasks. This is especially evident for happy voice stimuli, and it may be due to a difficulty introduced by splitting the explicit task into anger-neutral and happy-neutral blocks. Fourth, the use of selected discrete emotions does not allow to generalize for all existing emotions nor does it reveal the involvement of the basal ganglia and cerebellum in the processing of more complex emotions, such as pride, regret, jealousy, guilt, etc. Future research should explore these aspects to improve the generalizability and replicability of our findings. Finally, and though not the focus of the present article, the importance of acoustic parameters in these decisions should also be considered, especially voice fundamental frequency, pitch and spectral content, given their known role in vocal emotion processing (Banse & Scherer, 1996; Grandjean et al., 2006; Scherer, 1995).

### Implications and Future Directions

Our findings support the growing body of evidence highlighting the role of the cerebellum in emotion processing, extending its function beyond motor coordination to include predictive coding and integration of affective signals (Graybiel, 1998, 2008; Pierce & Péron, 2022). The differential involvement of cerebellar lobules depending on task demands and attentional focus suggests a hierarchical organization, with explicit processing engaging higher-order associative regions and implicit processing relying on more automatic, sensorimotor-driven mechanisms.

The observed direct connectivity between the cerebellum and basal ganglia also provides insights into how these structures interact during emotional prosody perception. The preferential connectivity during implicit processing suggests that these subcortical networks play a critical role in the rapid and automatic detection of emotional cues, a function that may be particularly relevant for social communication and affective decision-making in real-world contexts.

Future research should explore how these mechanisms are affected in clinical populations with cerebellar or basal ganglia dysfunctions, such as in patients with Parkinson’s disease, cerebellar or basal ganglia stroke. Additionally, longitudinal studies investigating the impact of aging on these networks could provide further insights into how attentional focus modulates emotional prosody processing across the lifespan.

### Conclusion

Overall, our study demonstrates that attentional focus significantly modulates cerebellar and basal ganglia involvement in emotional prosody processing. Explicit emotion recognition recruits a broader network, including higher-order cerebellar and basal ganglia structures, while implicitly processing voice emotion relies more on automatic, sensorimotor mechanisms. These findings contribute to deepening our understanding of emotional prosody decoding in the brain and highlight the crucial importance of considering both attentional and emotional factors when investigating vocal emotion perception.

## Data and code availability

Data and codes will be made available through a unique DOI or URL on the free, open repository YARETA, which meets the FAIR guidelines, upon acceptance of the manuscript for publication.

## Funding

The present research was supported by Swiss National Science Foundation (SNSF) grants to JAP (PI) and FA (Co-PI) within the project “Influence of top-down mechanisms on cerebellar activity during vocal emotion decoding” (2023-2026), Grant N°: 105314_215015 through the University of Geneva. LS was supported by Alzheimer’s Association research grant (AACSF-22-922907). The funders had no role in data collection, discussion of content, preparation of the manuscript, or decision to publish.

## Acknowledgments

We warmly thank the volunteers and the team of the Brain and Behavioral Laboratory of the University of Geneva, Switzerland, for their help with setting-up MRI sequences and with MRI data acquisition.

## Author contributions

LC and MT contributed equally by writing the first draft of the manuscript together, analyzing and interpreting the data and creating the figures; MT additionally acquired the data. IMC and EC acquired the data and helped analyze the data. JP designed and programmed the access to the study. AC helped with data analysis. DG, FA and JP supervised the study and allowed for an access to the imaging laboratory and funding; JP additionally designed the paradigm. All authors edited the manuscript.

## Ethics

This study was conducted following the written informed consent of all participants (N=28), and according to the strict regulations of the University of Geneva and the declaration of Helsinki. Participants were informed they could exit the study without justification any time they wanted or felt it was necessary for them. The state-wise ethics committee of Geneva University Hospital (Comité Cantonal d’Ethique en Recherche, CCER) reviewed and accepted our ethics application (number is 2024-00174).

## Notes

Disclosure: The authors report no conflicts of interest whatsoever.

### Competing Interest Statement

The authors have declared no competing interest.

### Summary of Updates

Revised version with considerable changes following reviews/revisions.

