## Supplementary figures and tables for "Influence of attention mechanisms on cerebellar and basal ganglia activity during vocal emotion decoding"

Supplementary material

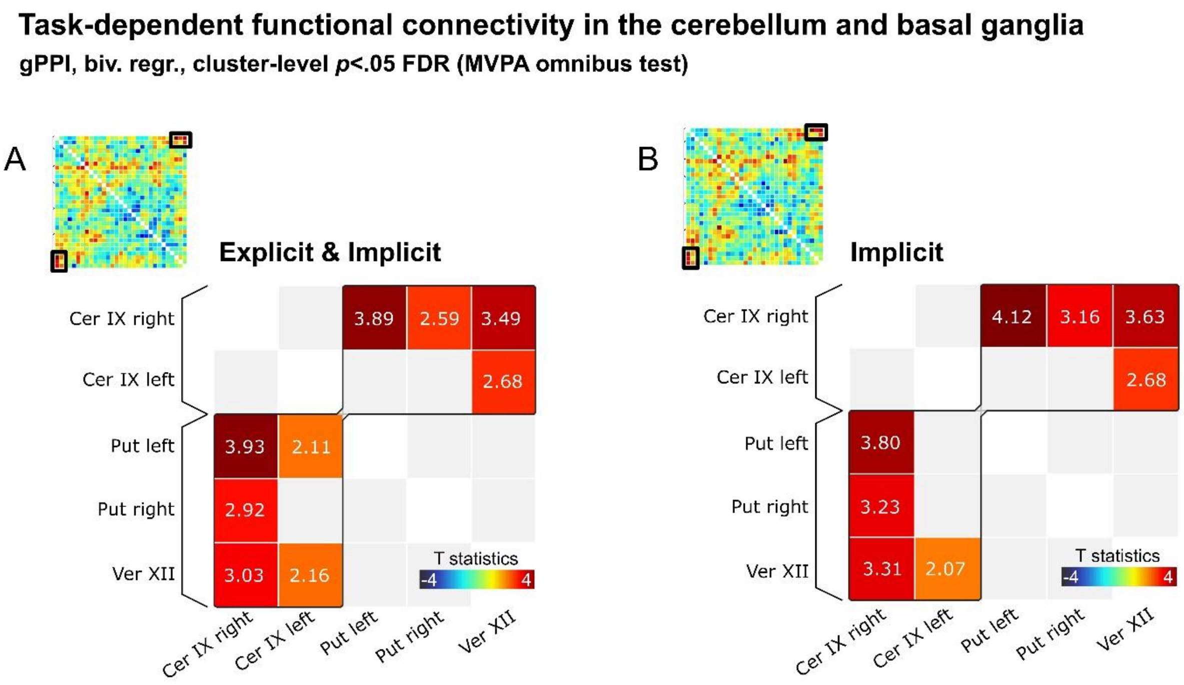


**Fig.S1: Functional connectivity for both and each task in the basal ganglia and cerebellum.** The level of basal ganglia and cerebellum task-based functional connectivity was computed by creating an interaction map, for each ROI as seed, of the main psychological (task) and physiological (the ROI) term, namely a psychophysiological interaction (PPI) with all ROIs taken at once and used as seed and target (generalized PPI or gPPI) and using bivariate regression. This analysis was computed for both tasks taken together ('A') and for each task, with results surviving thresholding only for the implicit task ('B'). Results are thresholded using parametric multivariate statistics and cluster-level inference with MVPA omnibus test and *p*<.05 FDR correction and two-tailed T-statistics. Cer: cerebellar lobule; Put: putamen; Ver: vermis.

**Explicit vs Implicit vocal emotion processing**
**p<.05 FDR, k>10**

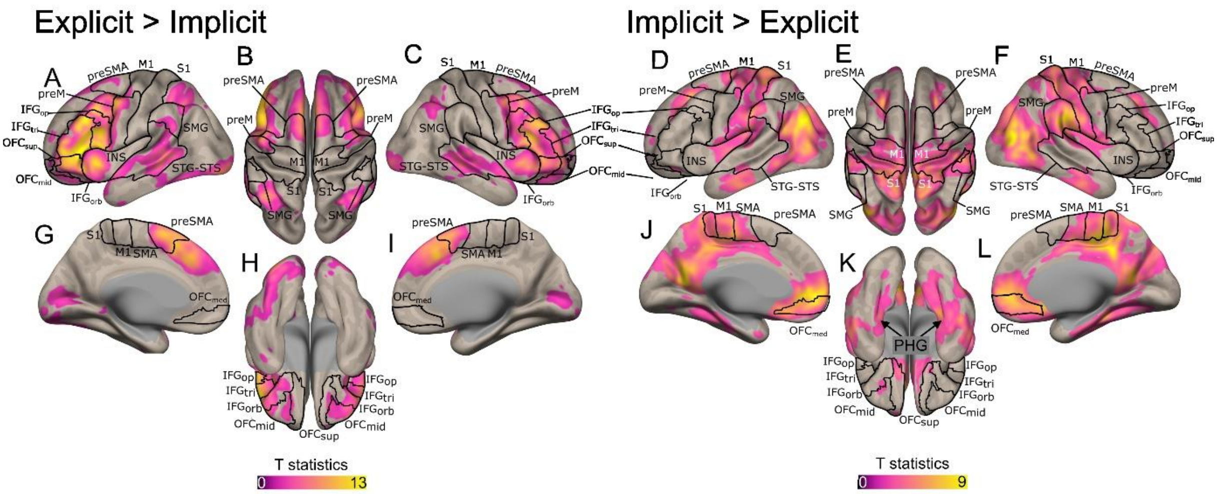


**Fig.S2: Whole brain activations contrasting the levels of the Task factor.** Lateral and medial cortical as well as basal ganglia and cerebellar activations were identified when contrasting explicit to implicit emotion processing tasks (Explicit>Implicit: lateral cortex 'A,B,C,H'; medial cortex 'G,I'; Implicit>Explicit: lateral cortex 'D,E,F,K'; medial cortex 'J,L'). All activations reported using an FDR corrected voxel-wise threshold of $p<.05$. Color bars represent T-values. IFG: inferior frontal gyrus; INS: insula; STG: superior temporal gyrus; STS: superior temporal sulcus; preSMA: pre-supplementary motor area; SMA: supplementary motor area; preM: premotor cortex; M1: primary motor cortex; S1: primary somatosensory cortex; SMG: supramarginal gyrus. Caud: caudate nucleus; Put: putamen; Thal: thalamus; OFC: orbitofrontal cortex; Cer: cerebellar lobule; Hipp: hippocampus. Suffixes: orb, *pars orbitalis*; tri, *pars triangularis*; op, *pars opercularis*; sup, superior; mid, middle; med, medial.

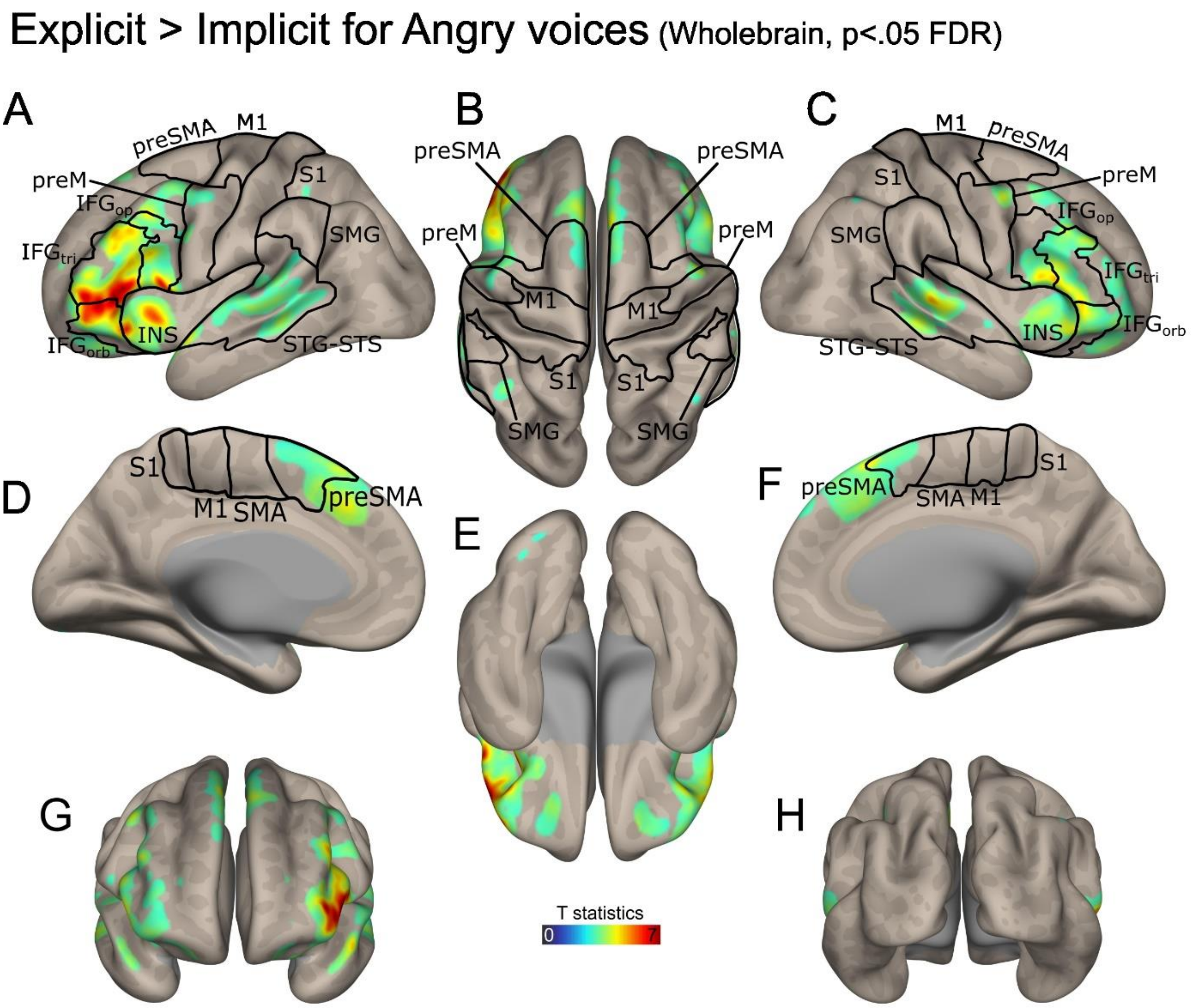


**Fig.S3: Whole brain activations contrasting Explicit > Implicit for angry voices.** Lateral and medial cortical activations were highlighted (lateral cortex ‘A,B,C,G,H’; medial cortex ‘D,E,F). All activations reported using an FDR corrected voxel-wise threshold of $p<.05$. Color bars represent T-values. IFG: inferior frontal gyrus; INS: insula; STG: superior temporal gyrus; STS: superior temporal sulcus; preSMA: pre-supplementary motor area; SMA: supplementary motor area; preM: premotor cortex; M1: primary motor cortex; S1: primary somatosensory cortex; vmPFC: ventromedial prefrontal cortex; SMG: supramarginal gyrus. Suffixes: tri, pars triangularis; op, pars opercularis.

### Explicit > Implicit for Happy voices (Wholebrain, p<.05 FDR)

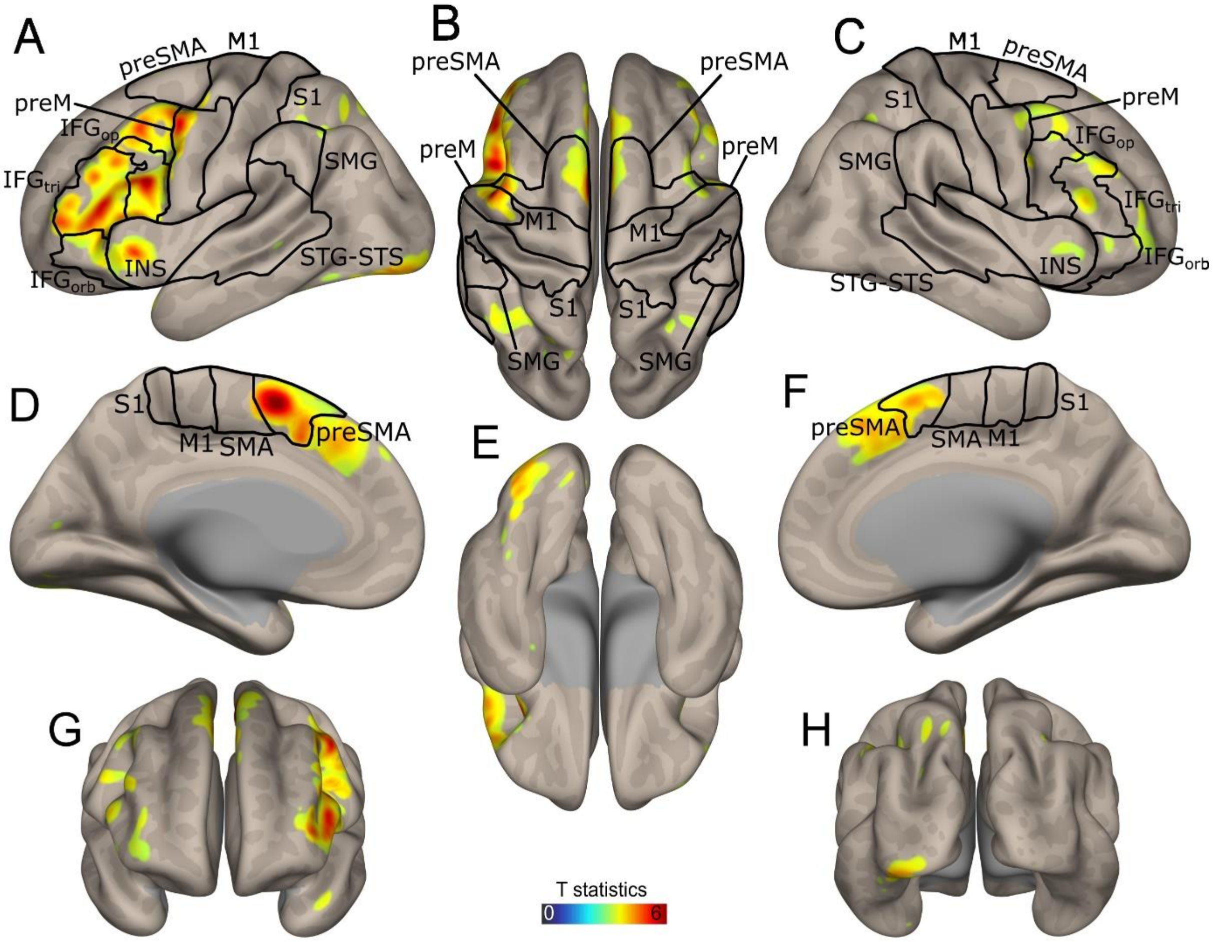


**Fig.S4: Whole brain activations contrasting Explicit > Implicit for happy voices.** Lateral and medial cortical activations were highlighted (lateral cortex 'A,B,C,G,H'; medial cortex 'D,E,F). All activations reported using an FDR corrected voxel-wise threshold of $p<.05$. Color bars represent T-values. IFG: inferior frontal gyrus; INS: insula; STG: superior temporal gyrus; STS: superior temporal sulcus; preSMA: pre-supplementary motor area; SMA: supplementary motor area; preM: premotor cortex; M1: primary motor cortex; S1: primary somatosensory cortex; vmPFC: ventromedial prefrontal cortex; SMG: supramarginal gyrus. Suffixes: tri, pars triangularis; op, pars opercularis.

Explicit > Implicit for Neutral voices (Wholebrain, p<.05 FDR)

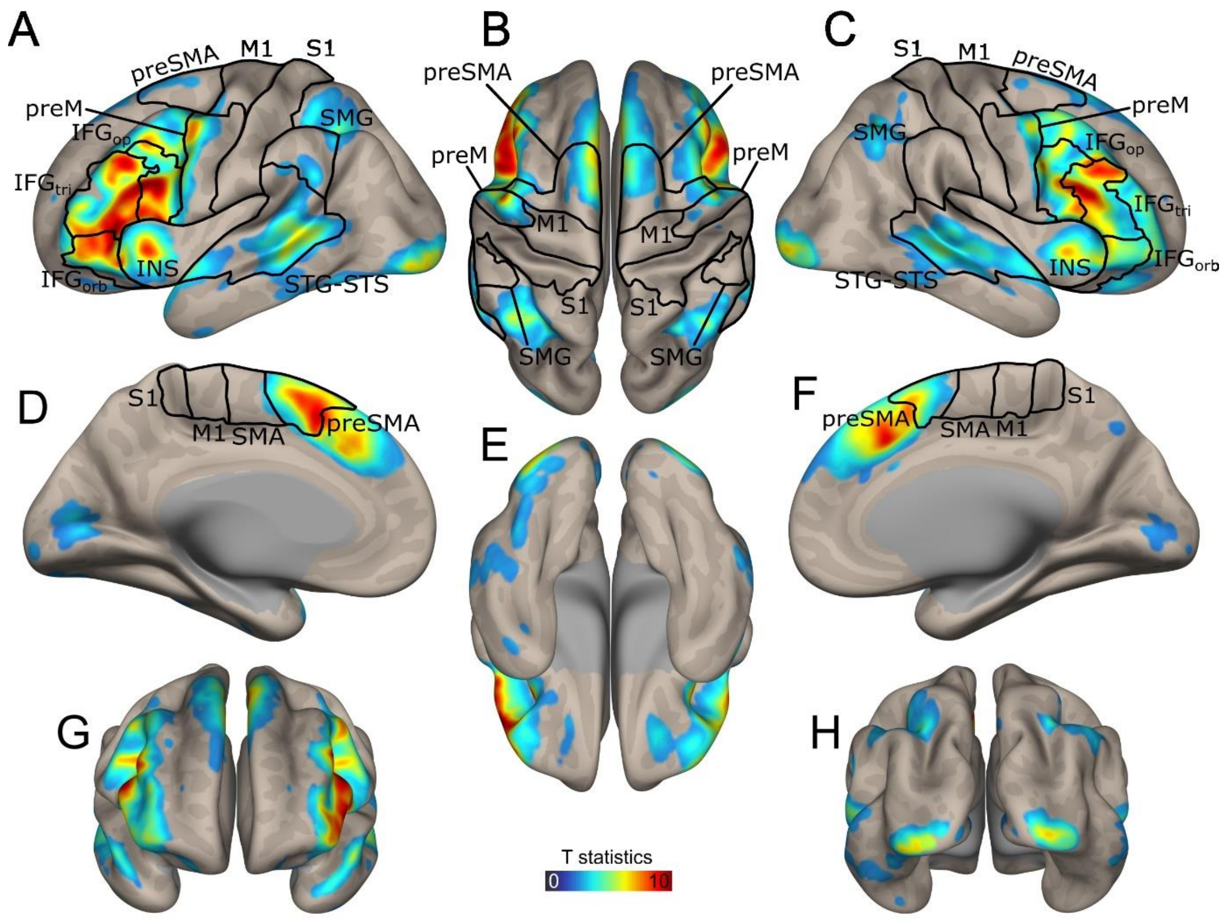


**Fig.S5: Whole brain activations contrasting Explicit > Implicit for neutral voices.** Lateral and medial cortical activations were highlighted (lateral cortex 'A,B,C,G,H'; medial cortex 'D,E,F). All activations reported using an FDR corrected voxel-wise threshold of $p<.05$. Color bars represent T-values. IFG: inferior frontal gyrus; INS: insula; STG: superior temporal gyrus; STS: superior temporal sulcus; preSMA: pre-supplementary motor area; SMA: supplementary motor area; preM: premotor cortex; M1: primary motor cortex; S1: primary somatosensory cortex; vmPFC: ventromedial prefrontal cortex; SMG: supramarginal gyrus. Suffixes: tri, pars triangularis; op, pars opercularis.

Implicit > Explicit for Angry voices (Wholebrain, p<.05 FDR)

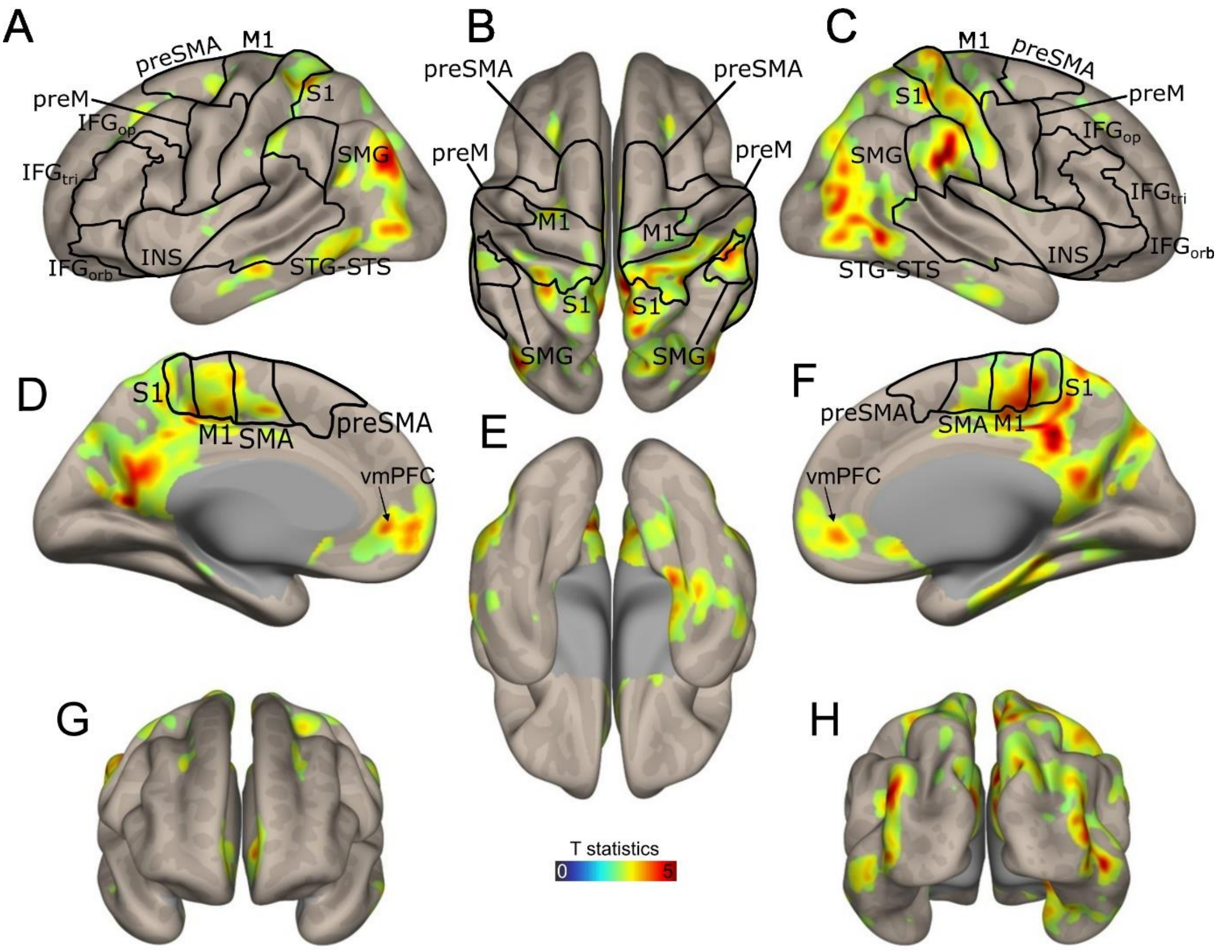


**Fig.S6: Whole brain activations contrasting Implicit > Explicit for angry voices.** Lateral and medial cortical activations were highlighted (lateral cortex 'A,B,C,G,H'; medial cortex 'D,E,F). All activations reported using an FDR corrected voxel-wise threshold of $p<.05$. Color bars represent T-values. IFG: inferior frontal gyrus; INS: insula; STG: superior temporal gyrus; STS: superior temporal sulcus; preSMA: pre-supplementary motor area; SMA: supplementary motor area; preM: premotor cortex; M1: primary motor cortex; S1: primary somatosensory cortex; vmPFC: ventromedial prefrontal cortex; SMG: supramarginal gyrus. Suffixes: tri, pars triangularis; op, pars opercularis.

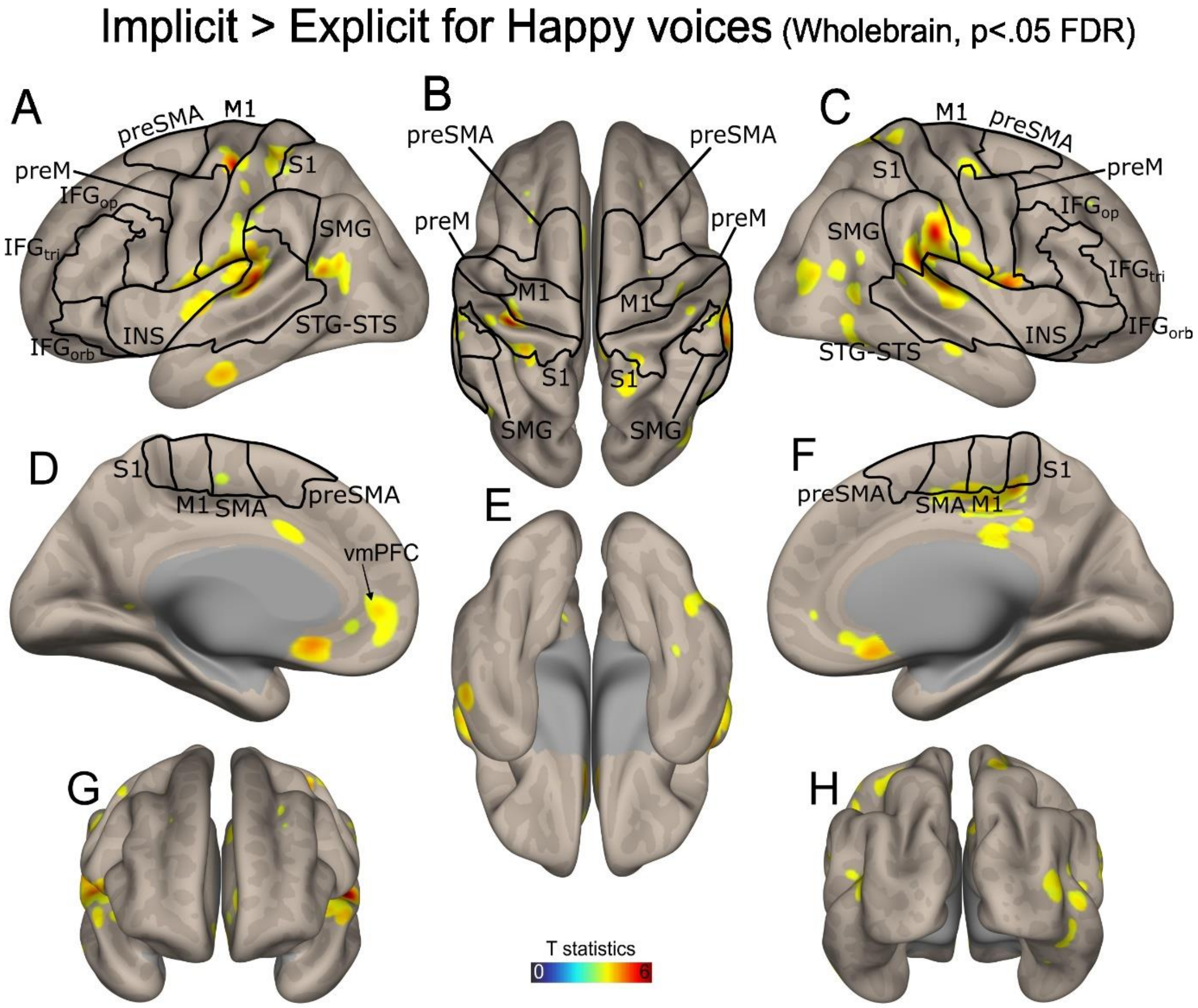


**Fig.S7: Whole brain activations contrasting Implicit > Explicit for happy voices.** Lateral and medial cortical activations were highlighted (lateral cortex 'A,B,C,G,H'; medial cortex 'D,E,F). All activations reported using an FDR corrected voxel-wise threshold of $p$<.05. Color bars represent T-values. IFG: inferior frontal gyrus; INS: insula; STG: superior temporal gyrus; STS: superior temporal sulcus; preSMA: pre-supplementary motor area; SMA: supplementary motor area; preM: premotor cortex; M1: primary motor cortex; S1: primary somatosensory cortex; vmPFC: ventromedial prefrontal cortex; SMG: supramarginal gyrus. Suffixes: tri, pars triangularis; op, pars opercularis.

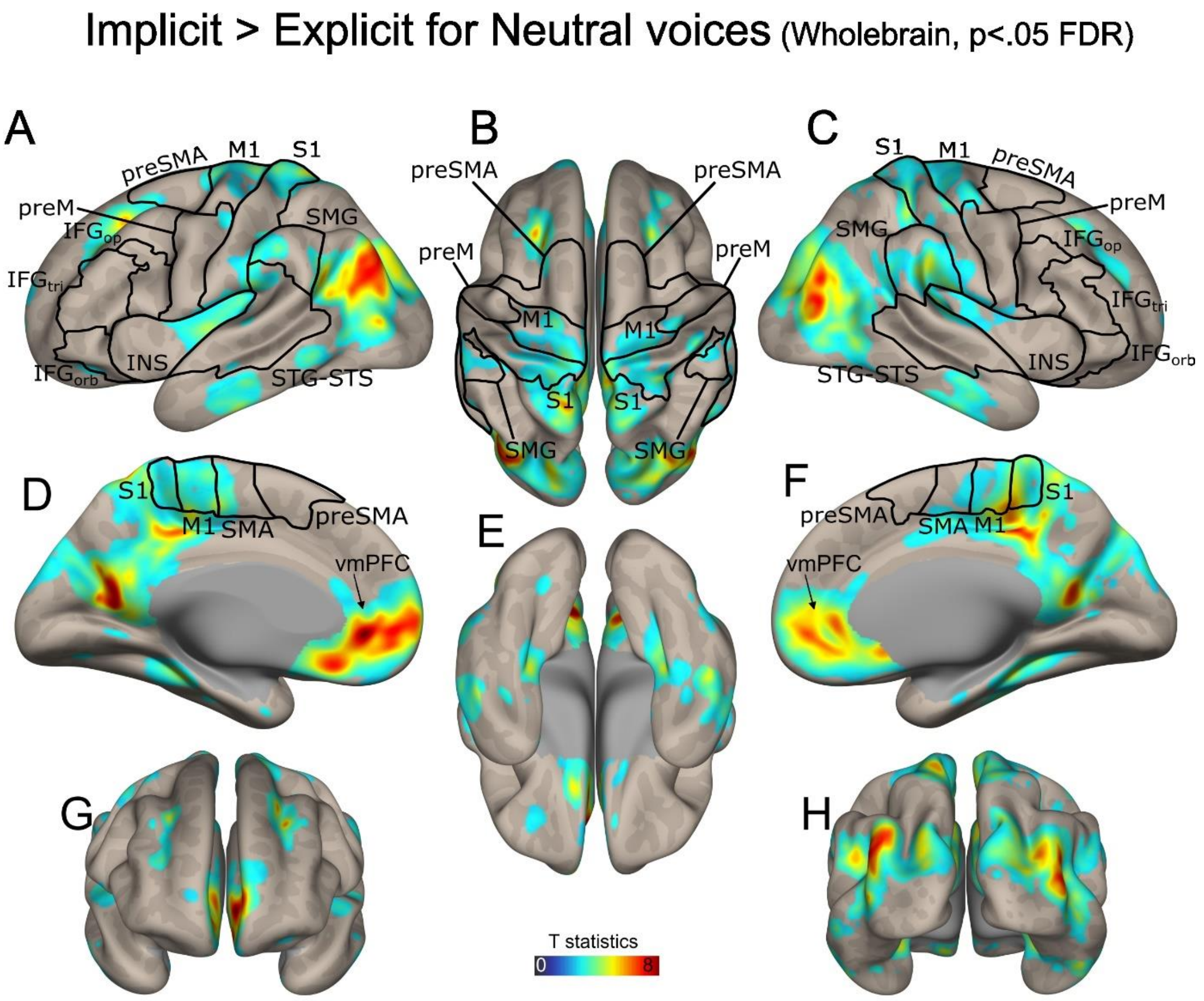


**Fig.S8: Whole brain activations contrasting Implicit > Explicit for neutral voices.** Lateral and medial cortical activations were highlighted (lateral cortex 'A,B,C,G,H'; medial cortex 'D,E,F). All activations reported using an FDR corrected voxel-wise threshold of $p<.05$. Color bars represent T-values. IFG: inferior frontal gyrus; INS: insula; STG: superior temporal gyrus; STS: superior temporal sulcus; preSMA: pre-supplementary motor area; SMA: supplementary motor area; preM: premotor cortex; M1: primary motor cortex; S1: primary somatosensory cortex; vmPFC: ventromedial prefrontal cortex; SMG: supramarginal gyrus. Suffixes: tri, pars triangularis; op, pars opercularis.

**Table S1: Descriptive statistics for behavioral data (Task and Emotion factors).**

| Response probability | | | |
|---|---|---|---|
| **Task** | **Voice emotion** | **Mean** | **Std** |
| **Explicit (Emotion)** | **Anger** | **0.95** | **0.22** |
| | **Happy** | **0.51** | **0.50** |
| | **Neutral** | **0.71** | **0.45** |
| **Implicit (Gender)** | **Anger** | **0.92** | **0.27** |
| | **Happy** | **0.98** | **0.14** |
| | **Neutral** | **0.98** | **0.13** |
| **Reaction times [msec]** | | | |
| **Task** | **Voice emotion** | **Mean** | **Std** |
| **Explicit (Emotion)** | **Anger** | **624.76** | **309.65** |
| | **Happy** | **769.79** | **385.51** |
| | **Neutral** | **612.80** | **311.98** |
| **Implicit (Gender)** | **Anger** | **593.40** | **282.87** |
| | **Happy** | **582.59** | **252.93** |
| | **Neutral** | **523.64** | **236.57** |

**Std: standard deviation.**

**Table S2: Brain coordinates of above-threshold cluster activations, Explicit vs Implicit contrast.**

| Cluster size | T value | MNI X | MNI Y | MNI Z | Region | Hemisphere |
|---|---|---|---|---|---|---|
| 9952 | 12,93 | -54 | 26 | 2 | IFGtri | L |
| | 12,21 | -50 | 36 | 2 | IFGtri | L |
| | 11,99 | -48 | 22 | -6 | IFGtri | L |
| | 11,65 | -32 | 22 | -4 | INSant | L |
| | 11,63 | -54 | 16 | 6 | IFGop | L |
| 14456 | 11,30 | -2 | 20 | 48 | preSMA | L |
| | 10,76 | 4 | 28 | 42 | ACC | R |
| | 10,33 | 48 | 40 | 20 | DLPFC | R |
| | 10,18 | -2 | 32 | 38 | ACC | L |
| | 10,09 | 52 | 28 | 20 | IFGop | R |
| 5586 | 7,82 | -10 | -78 | -28 | Cerebellum Crus I | L |
| | 7,31 | -34 | -86 | -12 | Cerebellum Crus I | L |
| | 6,70 | -20 | -96 | -8 | Cerebellum Crus II | L |
| | 6,65 | 12 | -78 | -30 | Cerebellum Crus I | R |
| | 5,74 | 38 | -64 | -28 | Cerebellum Crus II | R |
| 1641 | 7,56 | -38 | -50 | 44 | IPL | L |
| | 4,91 | -30 | -64 | 46 | IPL | L |
| 798 | 6,41 | 38 | -58 | 48 | IPL | R |
| 444 | 6,01 | 30 | -94 | -6 | LOC | R |
| 1790 | 5,63 | 12 | 8 | 8 | Caudate nucleus | R |
| | 5,36 | 14 | 2 | 14 | Caudate nucleus | R |
| | 4,78 | -12 | 10 | 0 | Caudate nucleus | L |
| | 4,68 | -10 | -6 | 2 | Thalamus | L |
| | 4,22 | 8 | -12 | -12 | SubstNigra | R |
| 589 | 4,76 | -6 | -82 | 6 | Calcarine | L |
| 157 | 4,11 | -22 | 54 | -12 | OFCsup | L |
| | 4,09 | -20 | 36 | -22 | OFCsup | L |
| | 3,99 | -20 | 44 | -18 | OFCsup | L |
| 22 | 3,30 | -40 | -4 | -40 | ITG | L |
| 44 | 3,19 | 62 | -48 | -20 | ITG | R |
| | 3,17 | 64 | -40 | -16 | ITG | R |
| 15 | 2,95 | -12 | -42 | -4 | Lingual | L |
| 16 | 2,88 | -2 | -30 | -2 | Brainstem | L |
| 14 | 2,79 | -24 | -36 | -46 | Cerebellum lobule X | L |

**IFG: inferior frontal gyrus; INS: insula; preSMA: pre supplementary motor area; ACC: anterior cingulate cortex; DLPFC: dorsolateral prefrontal cortex; IPL: inferior parietal lobule; LOC: lateral occipital cortex; SubstNigra: substantia nigra; OFCsup: orbitofrontal cortex superior part; ITG: inferior temporal gyrus. Suffixes: ant, anterior; tri: pars triangularis; op, pars opercularis.**

**Table S3: Brain coordinates of above-threshold cluster activations, Implicit vs Explicit contrast.**

| Cluster size | T value | MNI X | MNI Y | MNI Z | Region | Hemisphere |
|---|---|---|---|---|---|---|
| 6873 | 9,24 | -4 | 52 | -4 | vmPFC | L |
| | 7,34 | 4 | 22 | -10 | vmPFC | R |
| | 6,80 | 8 | 52 | -4 | vmPFC | R |
| | 6,42 | 6 | 40 | -6 | vmPFC | R |
| | 6,11 | 26 | 34 | 34 | DLPFC | R |
| 47528 | 8,90 | 10 | -32 | 44 | PCC | R |
| | 8,89 | 46 | -68 | 20 | MTGpost | R |
| | 8,55 | -42 | -78 | 30 | LOCmid | L |
| | 8,49 | -16 | -58 | 14 | PCC | L |
| | 8,14 | -6 | -60 | 20 | Precuneus | L |
| 1094 | 6,67 | -22 | 26 | 44 | DLPFC | L |
| | 6,56 | -26 | 32 | 40 | DLPFC | L |
| 265 | 3,90 | -28 | -48 | -54 | Cerebellum lobule VIII | L |
| | 3,68 | -12 | -54 | -48 | Cerebellum lobule IX | L |
| | 3,35 | -22 | -54 | -50 | Cerebellum lobule VIII | L |
| 65 | 3,87 | 20 | -50 | -52 | Cerebellum lobule VIII | R |

**vmPFC: ventromedial prefrontal cortex; DLPFC: dorsolateral prefrontal cortex ; PCC : posterior cingulate cortex; MTG: middle temporal gyrus; LOC: lateral occipital cortex. Suffixes : post, posterior ; mid, mid.**

**Table S4: Brain coordinates of above-threshold cluster activations, Task * Emotion factor (Explicit > Implicit for angry, happy, neutral voices separately).**

| | Cluster size | T value | MNI X | MNI Y | MNI Z | Region | Hemisphere |
|---|---|---|---|---|---|---|---|
| **Angry voice** | 4286 | 7,53 | -54 | 28 | 2 | IFGtri | L |
| | 2808 | 5,73 | 56 | 26 | -6 | IFGorb | R |
| | 769 | 5,34 | 64 | -28 | 2 | STGpost | R |
| | 1291 | 5,30 | 4 | 28 | 60 | preSMA | R |
| | 831 | 4,42 | -58 | -34 | 0 | STSmid | L |
| | 84 | 4,25 | -20 | 42 | -18 | OFCmed | L |
| | 61 | 3,97 | 22 | 48 | -16 | OFCmid | R |
| | 257 | 3,96 | -12 | -76 | -32 | Crus II | L |
| | 144 | 3,93 | 14 | 0 | 16 | Caudate | R |
| | 58 | 3,80 | -38 | -48 | 42 | IPL | L |
| | 42 | 3,66 | -38 | -30 | 8 | A1 | L |
| | 67 | 3,51 | -44 | -72 | -32 | Crus I | L |
| | 71 | 3,47 | 36 | 56 | 10 | MFG | R |
| | 39 | 3,43 | -16 | -96 | -8 | LOCinf | L |
| | 22 | 3,39 | 6 | 56 | 38 | SFGmed | R |
| **Happy voice** | 1340 | 6,03 | -4 | 6 | 60 | preSMA | L |
| | 2751 | 5,86 | -50 | 10 | 48 | preM | L |
| | 141 | 5,11 | -32 | 22 | -6 | INSant | L |
| | 233 | 4,98 | -36 | -84 | -16 | FFC | L |
| | 167 | 4,34 | 28 | -66 | -26 | Cer VI | R |
| | 303 | 4,23 | 54 | 14 | 36 | IFGop | R |
| | 219 | 4,11 | 48 | 44 | 22 | MFG | R |
| | 326 | 4,06 | 6 | -74 | -24 | Ver VI | R |
| | 132 | 3,85 | -38 | -68 | -26 | Crus I | L |
| | 159 | 3,83 | -44 | -48 | 50 | IPL | L |
| | 52 | 3,68 | -44 | 22 | -24 | STGant | L |
| | 69 | 3,65 | -24 | -70 | 52 | SPL | L |
| | 21 | 3,29 | 34 | -56 | 44 | AG | R |
| | 11 | 3,24 | 34 | 24 | -4 | INSant | R |
| | 16 | 3,23 | 28 | -92 | -10 | LOCinf | R |
| | 10 | 3,23 | -46 | -62 | -16 | FFC | L |
| **Neutral voice** | 7566 | 11,24 | -54 | 26 | 0 | IFGtri | L |
| | 4454 | 10,67 | -2 | 20 | 48 | SMA | L |
| | 8991 | 10,05 | 52 | 22 | 22 | IFGtri | R |
| | 465 | 7,36 | 32 | -94 | -4 | LOCinf | R |

| | | | | | | |
|---|---|---|---|---|---|---|
| 4110 | 7,36 | -34 | -90 | -10 | LOCinf | L |
| 1514 | 6,33 | -34 | -52 | 42 | IPL | L |
| 2676 | 6,01 | 12 | 4 | 10 | Caudate | R |
| 800 | 5,82 | 38 | -56 | 46 | AG | R |
| 1187 | 5,56 | 12 | -78 | -30 | Crus I | R |
| 115 | 5,00 | 20 | 42 | -16 | OFCmed | R |
| 246 | 4,26 | -6 | -78 | 8 | Calc | L |
| 87 | 4,01 | 64 | -40 | -16 | ITGpost | R |
| 22 | 3,27 | -40 | -4 | -40 | ITGant | L |
| 15 | 3,25 | -20 | 36 | -22 | OFCmed | L |
| 34 | 2,91 | 10 | -82 | 4 | Calc | R |
| 10 | 2,78 | 6 | -68 | 48 | Prec | R |

**IFG: inferior frontal gyrus; INS: insula; preSMA: pre supplementary motor area; IPL: inferior parietal lobule; LOC: lateral occipital cortex; OFC: orbitofrontal cortex; ITG: inferior temporal gyrus; LOC: lateral occipital cortex; AG: angular gyrus; Calc: calcarine; Prec: precuneus; FFC: fusiform cortex; SPL: superior parietal lobule; STG: superior temporal gyrus; preM: premotor cortex; Ver: Vermis; Cer: cerebellar lobule; MFG: middle frontal gyrus; A1: primary auditory cortex; STS: superior temporal sulcus; SFG: superior frontal gyrus. Suffixes: ant, anterior; post, posterior; tri: pars triangularis; op, pars opercularis; orb, pars orbitalis; sup, superior; inf, inferior; med, medial.**

**Table S5: Brain coordinates of above-threshold cluster activations, Task * Emotion factor (Implicit > Explicit for angry voices).**

| Cluster size | T value | MNI X | MNI Y | MNI Z | Region | Hemisphere |
|---|---|---|---|---|---|---|
| 900 | 5,92 | -4 | 52 | -4 | OFCmed | L |
| 13629 | 5,82 | 8 | -40 | 34 | PCC | R |
| | 5,71 | -16 | -58 | 14 | Prec | L |
| | 5,66 | 54 | -64 | -4 | ITGpost | R |
| | 5,59 | 52 | -28 | 32 | IPL | R |
| 1178 | 5,43 | -40 | -80 | 30 | LOCmid | L |
| 810 | 4,85 | 26 | -36 | -12 | PHG | R |
| 271 | 4,70 | 0 | 14 | -6 | OLFbulb | R |
| 129 | 4,30 | -64 | -20 | -16 | MTGmid | L |
| 83 | 3,93 | -40 | -56 | 26 | AG | L |
| 142 | 3,50 | 22 | -58 | -8 | FFC | R |
| 66 | 3,47 | -34 | -24 | -14 | Hipp | L |
| 106 | 3,45 | -64 | -24 | 34 | SMG | L |
| 49 | 3,31 | 34 | 6 | 12 | INS | R |
| 11 | 3,09 | -22 | -34 | -18 | FFC | L |
| 17 | 3,08 | 28 | -44 | -50 | Cer VIII | R |
| 23 | 3,03 | -34 | -2 | 12 | INS | L |

**OFC: orbitofrontal cortex; PCC: posterior cingulate cortex; Prec: precuneus; ITG: inferior temporal gyrus; IPL: inferior parietal lobule; LOC: lateral occipital cortex; PHG: parahippocampal gyrus; OLFbulb: olfactory bulb; MTG: middle temporal gyrus; AG: angular gyrus; FFC: fusiform cortex; Hipp: hippocampus; SMG: supramarginal gyrus; INS: insula; Cer: cerebellar lobule. Suffixes: post, posterior; med, medial.**

**Table S6: Brain coordinates of above-threshold cluster activations, Task * Emotion factor (Implicit > Explicit for happy voices).**

| Cluster size | T value | MNI X | MNI Y | MNI Z | Region | Hemisphere |
|---|---|---|---|---|---|---|
| 1175 | 6,22 | -52 | -30 | 10 | STGpost | L |
| | 4,24 | -50 | -24 | 24 | SMG | L |
| | 4,18 | -54 | -4 | 2 | STGmid | L |
| 297 | 5,44 | -36 | -26 | 54 | M1 | L |
| 1371 | 5,08 | 58 | -28 | 22 | SMG | R |
| | 4,99 | 54 | 4 | 4 | preM | R |
| 97 | 4,63 | 0 | 22 | -14 | OLFbulb | R |
| 439 | 4,45 | 12 | -34 | 52 | PCC | R |
| 182 | 4,43 | -42 | -58 | 16 | MTGpost | L |
| 136 | 4,14 | 46 | -50 | -2 | MTGpost | R |
| 62 | 4,11 | -52 | -18 | -24 | MTGmid | L |
| 76 | 4,06 | 20 | -48 | 62 | S1 | R |
| 92 | 3,77 | -12 | 48 | 2 | vmPFC | L |
| 130 | 3,76 | 44 | -66 | 20 | MTGpost | R |
| 29 | 3,61 | 44 | -12 | 56 | preM | R |
| 28 | 3,60 | 70 | -14 | -16 | MTGmid | R |
| 14 | 3,52 | 36 | -2 | 10 | INS | R |
| 10 | 3,39 | 10 | 32 | -12 | OFCmed | R |
| 10 | 3,37 | 40 | -40 | -16 | FFC | R |

**STG: superior temporal gyrus; SMG: supramarginal gyrus; M1: primary motor cortex; preM: premotor cortex; OLFbulb: olfactory bulb; OFC: orbitofrontal cortex; PCC: posterior cingulate cortex; S1: primary somatosensory cortex; vmPFC: ventromedial prefrontal cortex; MTG: middle temporal gyrus; FFC: fusiform cortex; INS: insula. Suffixes: post, posterior; med, medial.**

**Table S7: Brain coordinates of above-threshold cluster activations, Task * Emotion factor (Implicit > Explicit for neutral voices).**

| Cluster size | T value | MNI X | MNI Y | MNI Z | Region | Hemisphere |
|---|---|---|---|---|---|---|
| 29095 | 8,12 | -4 | 48 | -6 | OFCmed | L |
| | 7,60 | -14 | -58 | 16 | Prec | L |
| | 7,18 | -42 | -76 | 30 | LOCmid | L |
| | 6,99 | 12 | -32 | 42 | PCC | R |
| 887 | 6,10 | -24 | 30 | 42 | MFGmid | L |
| 410 | 4,99 | -34 | -32 | -14 | PHG | L |
| 473 | 4,96 | 52 | -28 | -26 | ITGmid | R |
| 1838 | 4,86 | -40 | -10 | 2 | INS | L |
| 367 | 4,72 | -58 | -20 | -28 | ITGmid | L |
| 49 | 4,59 | -30 | -48 | -54 | Cer VIII | L |
| 79 | 3,96 | -54 | -54 | -6 | ITGpost | L |
| 37 | 3,38 | 2 | -72 | 58 | Prec | R |
| 21 | 3,08 | 50 | -58 | -18 | ITGpost | R |
| 11 | 2,77 | 42 | -26 | 46 | S1 | R |
| 13 | 2,77 | 22 | -12 | -32 | PHG | R |

**OFC: orbitofrontal cortex; Prec: precuneus; LOC: lateral occipital cortex; PCC: posterior cingulate cortex; MFG: middle frontal gyrus; PHG: parahippocampal gyrus; ITG: inferior temporal gyrus; S1: primary somatosensory cortex; Cer: cerebellar lobule; INS: insula. Suffixes: post, posterior; med, medial.**

**Table S8: Brain coordinates of above-threshold cluster activations, Emotion factor (difference between happy and neutral voices), explicit task.**

| Cluster size | T value | MNI X | MNI Y | MNI Z | Region | Hemisphere |
|---|---|---|---|---|---|---|
| 1693 | 5,72 | -8 | -68 | 44 | Precuneus | L |
| | 5,68 | -24 | -74 | 30 | Precuneus | L |
| | 5,33 | -28 | -72 | 44 | Precuneus | L |
| | 5,10 | -12 | -64 | 30 | Precuneus | L |
| | 3,96 | -4 | -70 | 56 | Precuneus | L |
| 606 | 5,49 | 34 | -68 | 42 | LOC | R |
| 453 | 5,43 | -36 | -82 | -14 | FFC | L |
| 105 | 4,49 | -4 | -34 | 38 | PCC | L |
| 82 | 4,31 | 50 | -56 | -20 | FFC | R |
| 130 | 4,19 | 10 | -64 | 42 | Precuneus | R |
| | 4,16 | 8 | -72 | 46 | Precuneus | R |
| | 3,88 | 16 | -62 | 32 | Precuneus | R |

**LOC: lateral occipital cortex; FFC : fusiform cortex ; PCC : posterior cingulate cortex.**

**Table S9: Brain coordinates of above-threshold cluster activations, Emotion factor (difference between happy and neutral voices), implicit task.**

| Cluster size | T value | MNI X | MNI Y | MNI Z | Region | Hemisphere |
|---|---|---|---|---|---|---|
| 2062 | 7,68 | 38 | 6 | 32 | IFGop | R |
| | 6,60 | 60 | 30 | 4 | IFGtri | R |
| | 6,44 | 44 | 20 | 22 | IFGop | R |
| | 5,81 | 58 | 20 | 10 | IFGtri | R |
| | 5,74 | 48 | 6 | 50 | preSMA | R |
| 606 | 6,60 | 56 | 0 | -4 | STGmid | R |
| | 5,65 | 54 | -12 | 2 | STGpost | R |
| | 3,88 | 68 | -22 | 0 | STSpost | R |
| 288 | 6,48 | -36 | -84 | -10 | LOCinf | L |
| 1924 | 6,42 | -38 | 10 | 32 | IFGop | L |
| | 6,00 | -48 | 12 | 32 | IFGop | L |
| | 5,85 | -44 | 30 | 14 | IFGop | L |
| | 5,27 | -48 | 20 | 26 | IFGop | L |
| | 5,11 | -40 | 22 | 18 | IFGop | L |
| 365 | 5,72 | 0 | 14 | 60 | preSMA | Center |
| | 5,56 | -4 | 24 | 46 | preSMA | L |
| 230 | 5,64 | 36 | -86 | -2 | LOCinf | R |
| 32 | 5,58 | -36 | 20 | -30 | TP | L |
| 215 | 5,24 | -68 | -38 | 4 | STSpost | L |
| | 4,04 | -50 | -46 | 2 | STSpost | L |
| | 3,42 | -62 | -40 | 18 | STGpost | L |
| 249 | 4,83 | -4 | -70 | 42 | Precuneus | L |
| 165 | 4,36 | -30 | -60 | 40 | IPL | L |
| | 3,71 | -34 | -54 | 44 | IPL | L |
| 48 | 4,14 | 12 | 6 | 6 | Caudate nucleus | R |
| 31 | 4,05 | -46 | -62 | -18 | FFC | L |
| 55 | 4,02 | -30 | 22 | -2 | INSant | L |
| 64 | 4,02 | -14 | 2 | 12 | Caudate nucleus | L |
| 35 | 3,74 | -50 | -42 | -20 | ITGpost | L |
| | 3,62 | -60 | -44 | -18 | ITGpost | L |
| 16 | 3,72 | 2 | -14 | 4 | Thalamus | R |
| 61 | 3,62 | 34 | -58 | 50 | SPL | R |
| | 3,55 | 32 | -66 | 52 | SPL | R |
| 13 | 3,54 | -10 | -80 | -34 | Cerebellum Crus II | L |
| 35 | 3,52 | -30 | -64 | 58 | SPL | L |

**IFG: inferior frontal gyrus; preSMA: pre supplementary motor area; STG: superior temporal gyrus; STS: superior temporal sulcus; IPL: inferior parietal lobule; SPL: superior parietal lobule; FFC: fusiform cortex; TP: temporal pole; INS: insula; LOC: lateral occipital cortex; ITG: inferior temporal gyrus. Suffixes: ant, anterior; mid, mid; post, posterior; tri: pars triangularis; op, pars opercularis.**

**Table S10: Brain coordinates of above-threshold cluster activations, Emotion factor (difference between happy and angry voices), explicit task.**

| Cluster size | T value | MNI X | MNI Y | MNI Z | Region | Hemisphere |
|---|---|---|---|---|---|---|
| 6250 | 7,35 | -24 | -72 | 52 | Precuneus | L |
| | 6,62 | -6 | -68 | 44 | Precuneus | L |
| | 6,51 | -34 | -64 | 54 | SPL | L |
| | 6,13 | 10 | -74 | 48 | Precuneus | R |
| | 6,03 | 34 | -68 | 44 | SPL | R |
| 1098 | 5,44 | -36 | -82 | -16 | LOCinf | L |
| | 5,13 | -48 | -60 | -18 | FFC | L |
| | 4,97 | -60 | -46 | -16 | ITGpost | L |
| | 4,96 | -62 | -54 | -16 | ITGpost | L |
| | 4,74 | -46 | -70 | -12 | ITGpost | L |
| 808 | 5,13 | 46 | -76 | -8 | LOCinf | R |
| | 4,44 | 50 | -46 | -24 | ITGpost | R |
| | 3,86 | 60 | -40 | -12 | MTGpost | R |
| 851 | 4,76 | -48 | 10 | 48 | Precentral | L |
| | 4,10 | -46 | 14 | 30 | IFGop | L |
| | 3,92 | -42 | -2 | 52 | Precentral | L |
| | 3,78 | -42 | 6 | 38 | IFGop | L |
| 143 | 4,50 | 42 | 2 | 36 | IFGop | R |
| 84 | 4,17 | -2 | 8 | 58 | SMA | L |
| 256 | 4,03 | -46 | 44 | 18 | DLPFC | L |
| | 3,87 | -32 | 54 | 10 | DLPFC | L |
| | 3,60 | -46 | 34 | 14 | DLPFC | L |
| | 3,07 | -42 | 52 | 6 | DLPFC | L |
| 204 | 3,97 | 26 | -2 | 56 | SFG | R |
| | 3,58 | 32 | 8 | 56 | SFG | R |
| | 3,57 | 32 | -6 | 60 | SFG | R |
| | 3,32 | 38 | 4 | 60 | SFG | R |
| 45 | 3,88 | 38 | -30 | -24 | FFC | R |
| 36 | 3,87 | 20 | 4 | -4 | Putamen | R |
| 55 | 3,83 | -22 | -68 | -12 | FFC | L |
| 102 | 3,74 | 32 | -64 | -18 | Cerebellum lobule VI | R |
| 11 | 3,58 | 24 | 66 | 4 | FP | R |
| 27 | 3,53 | 24 | -30 | -18 | PHGpost | R |

**IFG: inferior frontal gyrus; SMA: supplementary motor area; IPL: inferior parietal lobule; SPL: superior parietal lobule; FFC: fusiform cortex; MTG: middle temporal gyrus; FP: frontal pole; INS: insula; LOCinf: inferior lateral occipital cortex; DLPFC: dorsolateral prefrontal cortex; SFG: superior frontal gyrus; PHG: parahippocampal gyrus; ITG: inferior temporal gyrus. Suffixes: post, posterior; op, pars opercularis.**

**Table S11: Brain coordinates of above-threshold cluster activations, Emotion factor (difference between angry and happy voices), explicit task.**

| Cluster size | T value | MNI X | MNI Y | MNI Z | Region | Hemisphere |
|---|---|---|---|---|---|---|
| 881 | 6,11 | -56 | -34 | 12 | STGpost | L |
| | 5,37 | -50 | -18 | 16 | preMotor | L |
| | 4,66 | -36 | -32 | 10 | STGpost | L |
| | 4,63 | -46 | -40 | 18 | STGpost | L |
| | 4,18 | -40 | -38 | 2 | STGpost | L |
| 225 | 5,01 | 54 | -36 | 16 | STGpost | R |
| | 4,54 | 62 | -26 | 6 | STGpost | R |
| 100 | 4,60 | -58 | -8 | 6 | STGmid | L |
| 78 | 4,57 | 54 | 8 | 6 | IFGop | R |
| 44 | 4,39 | 2 | -32 | -6 | Brainstem | R |
| 57 | 4,31 | 0 | 38 | 6 | ACC | Center |
| 26 | 4,27 | -40 | -10 | 2 | INSpost | L |
| 13 | 4,14 | 46 | -10 | -4 | PT | R |
| 27 | 4,10 | 12 | -58 | -46 | Cerebellum lobule IX | R |
| 22 | 4,08 | -6 | 24 | -14 | vmPFC | L |
| 28 | 4,08 | 8 | 50 | 4 | ACC | R |
| 22 | 3,98 | 50 | 6 | 18 | IFGop | R |
| 15 | 3,85 | -58 | 22 | 6 | IFGtri | L |

**STG: superior temporal gyrus; IFG: inferior frontal gyrus; ACC: anterior cingulate cortex; INS: insula; PT: planum temporale; vmPFC: ventromedial prefrontal cortex. Suffixes: mid, mid; post, posterior; tri: pars triangularis; op, pars opercularis.**

**Supplementary results**

*Reaction times data*

Reaction times data of each task were analyzed next. Linear regression was used as a function of the Task and Emotion factors (fixed effects)—including the split into two runs, and also taking into consideration relevant random effects (see Methods). Since the tasks are rather 'easy' to perform and the fact that no instruction concerning response time was given to the participants, reaction times data were of no interest in the present paper, but were examined anyway for completeness.

Reaction times were predicted by the Task with faster values for the gender task ($\chi^2(1)=19.96$, $p<.001$), voice Emotion ($\chi^2(2)=7.03$, $p<.05$), Run ($\chi^2(1)=23.06$, $p<.001$) and the interaction between Task and Emotion ($\chi^2(2)=17.53$, $p<.001$). Across tasks, angry and happy voices yielded slower reaction times compared to neutral voices for the Emotion factor ($\chi^2(1)=5.92$, $p<.05$). The interaction between Task and Emotion was again mainly explained by a partly inverted pattern of data between tasks, in which happy and neutral compared to angry voices led to slower reaction times in the explicit task while the inverse was observed in the gender task ([Task Gender > Task Emotion * Angry > Neutral + Happy voices]: $\chi^2(1)=16.53$, $p<.001$). Additionally, differences were observed between angry vs happy voices with slower reaction times for happy voices in the Emotion task ($\chi^2(1)=5.21$, $p<.05$) but not in the Gender task ($\chi^2(1)=0.94$, $p>.10$)—but differences were also observed between tasks (slower reaction times for angry vs happy in the Gender task as opposed to the inverse in the Emotion task ($\chi^2(1)=16.83$, $p<.001$). Reaction times were also slower for angry vs neutral voices the Gender task ($\chi^2(1)=7.01$, $p<.01$) but not in the Emotion task ($\chi^2(1)=0.26$, $p>.10$), and between tasks ($\chi^2(1)=9.46$, $p<.01$) as well as slower reaction times for happy vs neutral voices in the Emotion task only ($\chi^2(1)=8.40$, $p<.01$; Gender task: $\chi^2(1)=2.50$, $p>.10$; Gender vs Emotion task:

$\chi^2(1)=2.56$, $p<.10$). Variance explained by the fixed effects was 31.25% ($R^2c$) while the full model including random effects ($R^2c$) explained 48.33% of the variance of the reaction times data. See Fig.S1 for details and illustration.
